# *In vivo* chemical reprogramming is associated with a toxic accumulation of lipid droplets hindering rejuvenation

**DOI:** 10.1101/2025.06.25.661123

**Authors:** Wayne Mitchell, Cecília G. de Magalhães, Alexander Tyshkovskiy, Yushi Uchida, Ludger J.E. Goeminne, Takaharu Ichimura, Emery L. Ng, Joseph V. Bonventre, Vadim N. Gladyshev

## Abstract

Partial reprogramming has emerged as a promising strategy to reset the epigenetic landscape of aged cells towards more youthful profiles. Recent advancements have included the development of chemical reprogramming cocktails that can lower the epigenetic and transcriptomic age of cells and upregulate mitochondrial biogenesis and oxidative phosphorylation. However, the ability for these cocktails to affect biological age in a mammalian aging model has yet to be tested. Here, we have analyzed the effects of partial chemical reprogramming on mitochondrial structure in aged mouse fibroblasts and tested its *in vivo* efficacy in genetically diverse male UM-HET3 mice. This approach increases the size of mitochondria, alters cristae morphology, causes an increased fusing of mitochondrial networks, and speeds up movement velocity. We also discover that partial chemical reprogramming upregulates the formation of intracellular lipid droplets. At lower doses, the chemical reprogramming cocktail can be safely administered to middle-aged mice using implantable osmotic pumps, albeit with no effect on the transcriptomic age of kidney or liver tissues, and only a modest effect on the expression of OXPHOS complexes. However, at higher doses, the cocktail causes a drastic reduction in body weight and body condition scores. In the livers and kidneys of these animals, we observe significant increases in oil red o staining indicative of excessive lipid droplet accumulation in these organs. Thus, the upregulation of lipid droplet formation during partial chemical reprogramming may cause toxicity hindering the rejuvenation of cells and tissues in aged mammals.

## INTRODUCTION

Aging is characterized by underlying molecular changes that manifest as phenotypic declines in function and increased risk of mortality (1). Aging displays considerable heterogeneity between different cell types, tissues, species, sexes, and even individual organisms of the same species and sex (2-6). However, declines in the functioning of essential cellular processes such as proteostasis, mitochondrial bioenergetic function, and DNA repair and structural maintenance are often shared across these groups, colloquially forming the hallmarks of aging. While several molecular theories of aging have been proposed, such as epigenetic information loss (7), deleteriome (8-10), antagonistic pleiotropy (11), or hyperfunction (12, 13), causal data that ties the therapeutic targeting of any of these biological features in question to a robust extension of mammalian lifespan is lacking. Given that directly intervening in the aging process could delay the onset of multiple aging-related diseases in humans simultaneously (14), there is a need to develop therapies that target aging despite our mechanistic insight shortcomings. One avenue that circumvents this issue is to focus on pharmacological, genetic, and dietary interventions that reduce biological age, which is more representative of health status and future risk of disease than chronological age (15-17). More advanced transcriptomic biomarkers (18, 19), epigenetic clocks (20-23), and proteomics predictors of mortality (24, 25) are continually being developed that can accurately and unbiasedly assess the impact of an intervention on mammalian aging and lifespan (20, 26).

The discovery of the Yamanaka factors (27) has led to considerable interest in adapting the forced expression of *Oct4*, *Sox2*, *Klf4*, and *Myc* (OSKM) to rejuvenate aged cells while simultaneously preserving their cellular identity (28, 29). Cyclic partial reprogramming with OSKM can extend the lifespan of progeria mice (30) and ameliorate some molecular aspects of aging in wild-type C57BL/6J mice (31), but it can also lead to teratoma formation and toxicity in the liver and intestines due to these specific cell types reprogramming faster than others (32, 33). However, partial reprogramming with only OSK has been shown to avoid these toxicity challenges, whilst still being able to restore vision in mouse models of glaucoma and aging and extend lifespan in wild-type mice (34, 35). Further efforts to translate OSK partial reprogramming to other model organisms and diseases are ongoing (36, 37). One challenge that persists in developing partial reprogramming as an effective gene therapy is the Adeno-associated virus (AAV) delivery of exogenous OSK to various organs. For example, while AAV9 can effectively induce OSK expression in the heart and liver, it is unable to do so in the brain or pancreas (35). Thus, there is an unmet need to either develop AAVs capable of delivering genes to the vast majority of organs, or to develop alternative strategies that can partially reprogram cells and tissues that do not rely on AAV delivery.

Recent advancements have also led to the full reprogramming of human somatic cells to iPSCs using multi-stage chemical cocktail treatments (38-40). iPSCs generated through chemical reprogramming have further been differentiated to islet-like cells and autologously transplanted into a patient with Type 1 diabetes, resulting in sustainably improved glycemic control (41). Short-term treatment, or partial chemical reprogramming, of fibroblasts with a subset of these compounds, in contrast, does not lead to de-differentiation but instead can affect several molecular features associated with aging (42). Furthermore, treatment with several of these cocktails can lower the transcriptomic age of both non-senescent and senescent mouse and human fibroblasts, respectively (43, 44). Mechanistically, partial chemical reprogramming is associated with strong upregulation of mitochondrial biogenesis and oxidative phosphorylation (OXPHOS) and global DNA hypomethylation that causes an additional reduction in cellular epigenetic age (43). However, it is still currently unknown if partial chemical reprogramming can affect biological age in other cell types and tissues, and the impact of partial chemical reprogramming on mitochondrial biology is not fully elucidated.

Therefore, to more fully characterize the mechanisms of partial chemical reprogramming, we analyzed the effects of 7c (repsox, tranylcypromine, valproate, forskolin, CHIR99021, DZNep, 4-[(E)-2-(5,6,7,8-Tetrahydro-5,5,8,8-tetramethyl-2-naphthalenyl)-1-propenyl]benzoic acid (TTNPB)) treatment on mitochondrial morphology, interaction networks, and movement dynamics using microscopy. In tandem, we delivered the 7c cocktail to genetically diverse UM-HET3 male mice to evaluate its ability to impact organismal biological age based on transcriptomic biomarkers in liver and kidney tissues. While lower doses were well-tolerated, they did not lead to robust gene expression changes in the livers and kidneys. In contrast, higher doses led to rapid weight loss and lowered body scores requiring euthanasia. Staining of the tissues from the animals that received high doses of the 7c cocktail revealed a strong increase in lipid droplet formation, whereas no other major histological defects were readily apparent in the livers and kidneys. Taken together, our results suggest that the increase in lipid droplet formation during 7c partial chemical reprogramming may lead to toxicity *in vivo*, hindering rejuvenation.

## MATERIALS AND METHODS

### Materials and animals

Cell culture media, consumables, and reagents were acquired from ThermoFisher Scientific (Waltham, MA). All other general chemicals and reagents were purchased from Sigma-Aldrich (St. Louis, MO). Male C57BL/6J mice were acquired from the National Institute on Aging (NIA) aged rodent colonies (Charles River Laboratories, Wilmington, MA). Male UM-HET3 mice were purchased from Charles River Laboratories. Mice were given standard 5053 diet (LabDiet, St. Louis, MO) and water *ad libitum*, were housed on a 12/12 hour light/dark cycle at ∼21°C, and were allowed to acclimate for at least 2 weeks prior to any experiments. C57BL/6J mice were housed in a specific-pathogen free (barrier) facility at 5 animals per cage, whereas UM-HET3 mice were housed in the in-and-out (non-barrier) facility at 3 animals per cage. All experiments using mice were performed in accordance with institutional guidelines for the use of laboratory animals and were approved by the Brigham and Women’s Hospital and Harvard Medical School Institutional Animal Care and Use Committees under Protocol # 2016N000368. Mouse ear fibroblasts were isolated from 25-month-old male C57BL/6J mice as previously described (43) and were used exclusively at low-passage numbers for all assays (< 5). Chemical reprogramming compounds were obtained from the vendors listed in Table S1 and were used at the concentrations listed. Analgesics and other surgery supplies for the osmotic pump implantations were acquired from Patterson Veterinary Supply (Loveland, CO).

### Embedding, sectioning, and TEM imaging

25-month-old mouse ear fibroblasts were treated with 7c or 50:50 PBS: DMSO vehicle for 6 days and cultured at 37°C, 5% CO_2_, and 3% O_2_ in DMEM/F12 supplemented with 10% FBS, 1X antibiotic and antimycotic, 1X non-essential amino acids, and 50 µM β-mercaptoethanol. Media and treatments were replaced once after 3 days. Cells were then seeded onto UV-sterilized Aclar plastic coverslips in 24-well plates and grown overnight at a starting cell density of 100,000 cells per well. After aspirating the media and rinsing once with PBS, cells were fixed in 2.5% glutaraldehyde, 1.25% paraformaldehyde, and 0.3% picric acid in 0.1M sodium cacodylate buffer (pH 7.4) for 1 hour at room temperature. Then, the cells were washed in 0.1M sodium cacodylate buffer (pH 7.4), post-fixed for 30 minutes in 1% osmium tetroxide and 1.5% potassium ferrocyanide, and washed twice in water and once in maleate buffer. Following this, cells were incubated in 1% uranyl acetate in maleate buffer for 30 minutes, followed by two washes with water and subsequent dehydration in grades of alcohol (5 minutes each: 50%, 70%, 95%, and 2 x 100%). Finally, the samples were embedded in TAAB Epon (TAAB Laboratories Equipment Ltd, Berkshire, England) and polymerized at 60°C for 48 hours.

After polymerization, the Aclar plastic was peeled off, and a small area (∼1 mm) of the flat embedded cells were cut out with a razor blade and remounted on an Epon block. Ultrathin sections (∼80 nm) were cut on a Reichert Ultracut-S microtome, picked up onto copper grids stained with lead citrate, and examined on a TecnaiG^2^ Spirit BioTWIN. Images were taken using an AMT 2k CCD camera. Measurements of mitochondrial size were performed manually in FIJI (45).

### Oil red o staining

25-month-old mouse ear fibroblasts were treated with 2c (repsox, tranylcypromine), 7c, or 50:50 PBS: DMSO vehicle for 6 days and cultured at 37°C, 5% CO_2_, and 3% O_2_ in DMEM/F12 supplemented with 10% FBS, 1X antibiotic and antimycotic, 1X non-essential amino acids, and 50 µM β-mercaptoethanol. Media and treatments were replaced once after 3 days. Cells were then seeded in 24-well plates and grown overnight at a starting cell density of 100,000 cells per well. Oil red o stock solution was prepared by dissolving 60 mg in 20 ml of 100% isopropanol and incubating at room temperature for 30 minutes. A working solution of oil red o was prepared by mixing 6 ml of stock solution with 4 ml of water, incubating at room temperature for 10 minutes, and filter sterilizing through a 0.22 µm filter. After aspirating the media, the cells were washed twice with PBS prior to fixation in 4% paraformaldehyde for 30 minutes on a rotator at room temperature. Following this, the cells were rinsed once with PBS, incubated for 5 minutes in 60% isopropanol, and then stained with oil red o for 15 minutes. After rinsing five times with PBS, the cells were imaged on an EVOS microscope (ThermoFisher Scientific, Waltham, MA). Following image acquisition, the PBS was aspirated and replaced with 200 µl of 100% isopropanol. After mixing and incubating for 2 minutes at room temperature, the solutions were withdrawn and added to a clear 96-well plate, and the absorbances were read at 492 nm on a BioTek Synergy H1 microplate reader (Agilent Technologies, Lexington, MA).

### Osmotic pump implantations

Osmotic minipump models 1004 and 2004 were purchased from DURECT Corporation (Cupertino, CA). Male UM-HET3 mice were anesthetized with isoflurane using a precision vaporizer (SomnoFlo Suite, Kent Scientific) and primed pumps filled with either vehicle (50% v/v DMSO/PEG300) or 7c (valproate, DZNep, forskolin, TTNPB, tranylcypromine, repsox, CHIR99021) were randomly assigned and subcutaneously implanted for 28 days. To account for any potential cage effects, mice were housed individually after the procedure. Following 28 days of treatment, mice were euthanized by CO_2_ and cervical dislocation, after which blood and tissues were collected and either fixed in 4% paraformaldehyde overnight or flash-frozen in liquid nitrogen and stored at -80°C.

### Confocal microscopy

For live cell imaging, 50,000 cells were seeded overnight on 35 mm Ibidi dishes with #1.5 polymer coverslips and stained for 30 minutes with 50 nM TMRM, 100 nM MitoView Green, and 10 µg/ml Hoescht 33342 in live cell imaging media (fibroblast growth media with 4 mM L-glutamine and without phenol red). Cells were then imaged on a CSU-W1 Yokogawa spinning disk with a 50 µm pinhole disk and an Andor Zyla 4.2 Plus sCMOS monochrome camera built around an inverted Nikon Ti microscope equipped with a heated enclosure and CO_2_ control. Cells were imaged with either a 20X/0.75 air or 100X/1.45 oil objective. Fluorescence intensities were quantified in FIJI (45), and mitochondrial morphology networks were analyzed using the Mitochondria Analyzer (MitoAnalyzer) plug-in (46) on the TMRM images. Time-lapse mitochondrial morphology and dynamics were quantified using Mitometer (47).

For imaging of fixed cells, 20,000 cells per well were seeded overnight on Nunc Lab-Tek II 8-chambered #1.5 coverglass that were coated with 0.2% gelatin for at least 1 hour prior to cell seeding. Cells were then rinsed with PBS, fixed in 4% paraformaldehyde, and permeabilized with 0.3% Triton X-100, all at room temperature. After blocking with 5% BSA for 30 minutes, cells were stained with mouse anti-Tom20 primary antibody for 1 hour, followed by rabbit anti-mouse AlexaFluor 561 secondary antibody for 1 hour and 1 µg/ml DAPI for 10 minutes, all at room temperature before three final washes in PBS. Cells were imaged using a Zeiss Axio Observer Z1 point-scanning confocal equipped with a LSM980 scan head, two multi-akali PMTs, environmental enclosure, and a 63X/1.4 oil objective. 3D (z-stack) images with either 0.2 µm or 0.5 µm step sizes of mitochondrial morphology networks were analyzed using the MitoAnalyzer FIJI plug-in (46).

### Tissue sectioning and staining

Fresh tissues were rinsed several times in PBS before fixing in 4% paraformaldehyde overnight at 4°C with gentle rotation. Then, tissues were cut with a razor blade and either infiltrated with 30% sucrose overnight or embedded in paraffin and cut into 5 µm sections on a microtome. The sucrose-infiltrated tissues were then frozen in OCT blocks and cut into 20 µm sections on a cryostat.

Paraffin-embedded sections were stained either with hematoxylin and eosin, or Masson’s trichrome according to standard, published procedures. OCT-embedded sections were stained with oil red o and counterstained with hematoxylin. Stained tissues were imaged on an Axio Imager 2 (Zeiss, Oberkochen, Baden-Württemberg, Germany) using an EC Plan-Neofluar 10X/0.3 M27 objective and an Axiocam 105 color camera. The percent of tissue stained with oil red o for control and partially chemically reprogrammed tissues were determined by using the trainable weka segmentation FIJI plug-in (48). Cellular and nuclear structures of hematoxylin- and eosin-stained images were determined using QuPath (49).

Frozen kidney sections were hydrated in PBS and treated with 1% SDS in PBS for 5 minutes. After blocking with 3% BSA in PBS, the sections were incubated with primary antibodies for 1 hour at room temperature. Goat polyclonal anti–KIM-1 (R&D Systems, Minneapolis, MN) was used as the primary antibody. For co-staining, biotinylated Lotus tetragonolobus lectin (LTL; Vector Laboratories, Newark, CA) was applied to detect proximal tubule brush borders. The sections were then incubated with Cy3-conjugated anti-goat IgG as the secondary antibody and streptavidin-DyLight 488 for 30 minutes at room temperature. For phalloidin staining, the sections were incubated with FITC-conjugated phalloidin (ThermoFisher Scientific, Waltham, MA) for 30 minutes at room temperature. After final washes, the slides were mounted using Vectashield mounting medium containing DAPI (Vector Laboratories, Newark, CA), and images were acquired using a Leica Stellaris 5 confocal microscope (Leica Microsystems, Wetzlar, Germany). Phalloidin-stained tissues were imaged less than a week after staining in a single imaging session.

### RNA-seq

RNA from ∼30 mg of flash-frozen tissue was purified using a direct-zol RNA miniprep kit (Zymo Research, Irvine, CA) after tissue homogenization using a Bead Ruptor Elite homogenizer (OMNI International, Kennesaw, GA) and 2 ml tubes pre-filled with 2.8 mm ceramic beads. RNA concentration was measured by Qubit using the RNA HS assay kit, and library prep and sequencing was performed as described previously (50). Reads were mapped to mouse genome (GRCm39) with STAR (version 2.7.11b) and counted via featureCounts (version 2.0.6). To filter out non-expressed genes, we left only genes with at least 10 reads in at least 50% of samples separately for liver and kidney datasets. Differentially expressed genes were identified separately for each tissue using one-way ANOVA model through edgeR package (51).

### Transcriptomic signature analysis

To explore how transcriptomic alterations induced by 7c treatment relate to known molecular signatures of aging, lifespan regulation, and reprogramming, we conducted functional enrichment analysis. We utilized reference datasets encompassing tissue-specific aging signatures from liver, kidney, and brain, as well as multi-tissue biomarkers of biological age and expected mortality, adjusted for chronological age (50, 52). Additionally, we included a hepatic signature of expected maximum lifespan in rodents (19), along with transcriptional profiles of iPSC reprogramming via OSKM factor induction in mice and across species (53). For each tissue (liver and kidney), we ranked genes using a signed log-transformed p-value metric estimated through differential expression analysis:

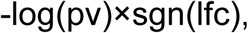

where pv and lfc are p-value and logFC of a certain gene, respectively, and sgn is the signum function (equal to 1, -1 and 0 if value is positive, negative or equal to 0, respectively). Ranked gene lists were then subjected to gene set enrichment analysis (GSEA) using the fgsea package in R, with 10,000 permutations and multilevel Monte Carlo sampling. Gene sets were drawn from the HALLMARK, KEGG, and REACTOME collections of the Molecular Signatures Database (MSigDB). The same enrichment pipeline was applied to reference signatures of aging, mortality, maximum lifespan, and iPSC reprogramming. To quantify similarities between 7c-induced and reference gene expression biomarkers, we computed Spearman correlations between normalized enrichment scores (NES).

### Transcriptomic age analysis

The filtered RNA-seq data were processed with Relative Log Expression (RLE) normalization, log-transformation, and scaling. Missing expression values for clock genes not detected in the dataset were imputed using their corresponding precomputed average values. Normalized gene expression profiles were centered to the median profile of control samples. Transcriptomic age (tAge) foe each sample was estimated using Bayesian Ridge multi-tissue transcriptomic clocks of chronological age and expected mortality (19). Module-specific transcriptomic clocks of chronological age were applied to scaled relative gene expression profiles of control, 2c-, and 7c-treated cells identified previously (43) using the same framework.

### Proteomics and phosphoproteomics analysis

To compare differential protein abundance and differential phosphorylation of 2c vs. 7c, we re-analyzed our TMT proteomics and phosphoproteomics datasets described previously (43), with the same msqrob2 (54, 55) model structure that includes the treatment (control, 2c, or 7c), age group (4 months, or 20 months), and the treatment:age group interaction as covariates. We also reused the GSEA described previously (43). For the targeted phosphoprotein enrichment analysis in Fig. 5C, we performed GSEA on the protein-level enrichments from the phosphopeptides (43) with only the four gene sets: “Structural constituent of muscle (Gene Ontology identifier (GO): 0008307)”, “Lipolysis in adipocytes (Kyoto Encyclopedia of Genes and Genomes identifier (KEGG): mmu04923)”, “Mitochondrion (GO: 0005739)”, and “mRNA splicing, via spliceosome (GO: 0000398)”.

### Western blotting

∼30 mg pieces of flash-frozen liver tissues were homogenized in 60 mM Tris-HCl, pH 6.8, 5% SDS buffer in 2 ml tubes pre-filled with 2.8 mm ceramic beads on a Bead Ruptor Elite homogenizer (OMNI International, Kennesaw, GA). Following homogenization, samples were spun at 12,000 x g for 2 minutes and the protein concentration of the supernatants was measured by BCA assay using a BioTek Synergy H1 microplate reader (Agilent Technologies, Lexington, MA). 15 µg of protein were loaded onto TGX 4-15% gradient gels (Bio-Rad, Hercules, CA) and run at 120V for ∼1 hour, transferred to PVDF membranes, and blotted for β-actin (Santa Cruz Biotechnology, Dallas, TX) or OXPHOS subunits (Thermo-Fisher, Waltham, MA). Blots were visualized by staining with IRDye 680RD secondary antibody and imaging on a LI-COR Odyssey Fc molecular imager (Lincoln, NE).

### Statistical analysis

For the proteomics and phosphoproteomics analysis, we used the msqrob2 and GSEA analyses described before (43). For comparison of transcriptomic ages (tAges), predicted by composite clocks, mixed-effects ANOVA model was implemented via the rma.uni function form the metafor package in R. For comparison of tAges estimated by module-specific clocks, one-way ANOVA was used. For omics analyses, p-values were adjusted for multiple testing with the Benjamini-Hochberg method (56). In other cases, p-values for comparisons between control and partially reprogrammed groups were determined by two-tailed unpaired t-tests assuming equal variance. Statistical tests were performed using either the R package limma (57) or GraphPad Prism version 10.0.0. All measurements consist of at least n = 3 independent biological replicates with their corresponding means (bar height) and standard deviations (error bars) depicted in the figures.

## RESULTS

Previously, we reported that partial chemical reprogramming with only the 7c and not the 2c (repsox, tranylcypromine) cocktail drove a significant increase in OXPHOS expression and mitochondrial spare respiratory capacity (43). Therefore, we further investigated the effects of 7c treatment on mouse fibroblast mitochondrial function, morphology, and interaction networks using live-cell imaging. Co-staining fibroblasts with mitochondrial transmembrane potential-independent (MitoView Green) and -dependent (TMRM) dyes revealed that 7c increased the fluorescence intensity of both compared to control cells treated with vehicle (**Fig. 1A**, left panels). Moreover, we used the TMRM signal to construct 2D masks of the mitochondrial networks (**Fig. 1A**, right panels). When we normalized the TMRM signal to account for changes in mitochondrial mass detected by MitoView Green, we still observed a significant increase in the mitochondrial transmembrane potential following 6 days of 7c treatment (**Fig. 1B**). We then used the 2D masks to measure the effects of partial chemical reprogramming with 7c on mitochondrial size and interaction networks. Following 7c treatment, we observed significant increases in mitochondrial area and perimeter, in addition to greater total branch length, number of branches, and branch junctions (**Fig. 1C**). We further collected time-lapse TMRM images (10 second intervals, 3 minutes total) and analyzed them to determine the impact of partial chemical reprogramming on mitochondrial movement dynamics. Relative to control cells, we observed that 7c treatment increased the average total distance traveled (**Fig. 2A**) and displacement (distance from starting position) (**Fig. 2B**) of the mitochondria, which was further supported by significant increases in mitochondrial movement speed (rate of movement) (**Fig. 2C**) and velocity (rate of movement in a given direction) (**Fig. 2D**).

**Figure 1:**
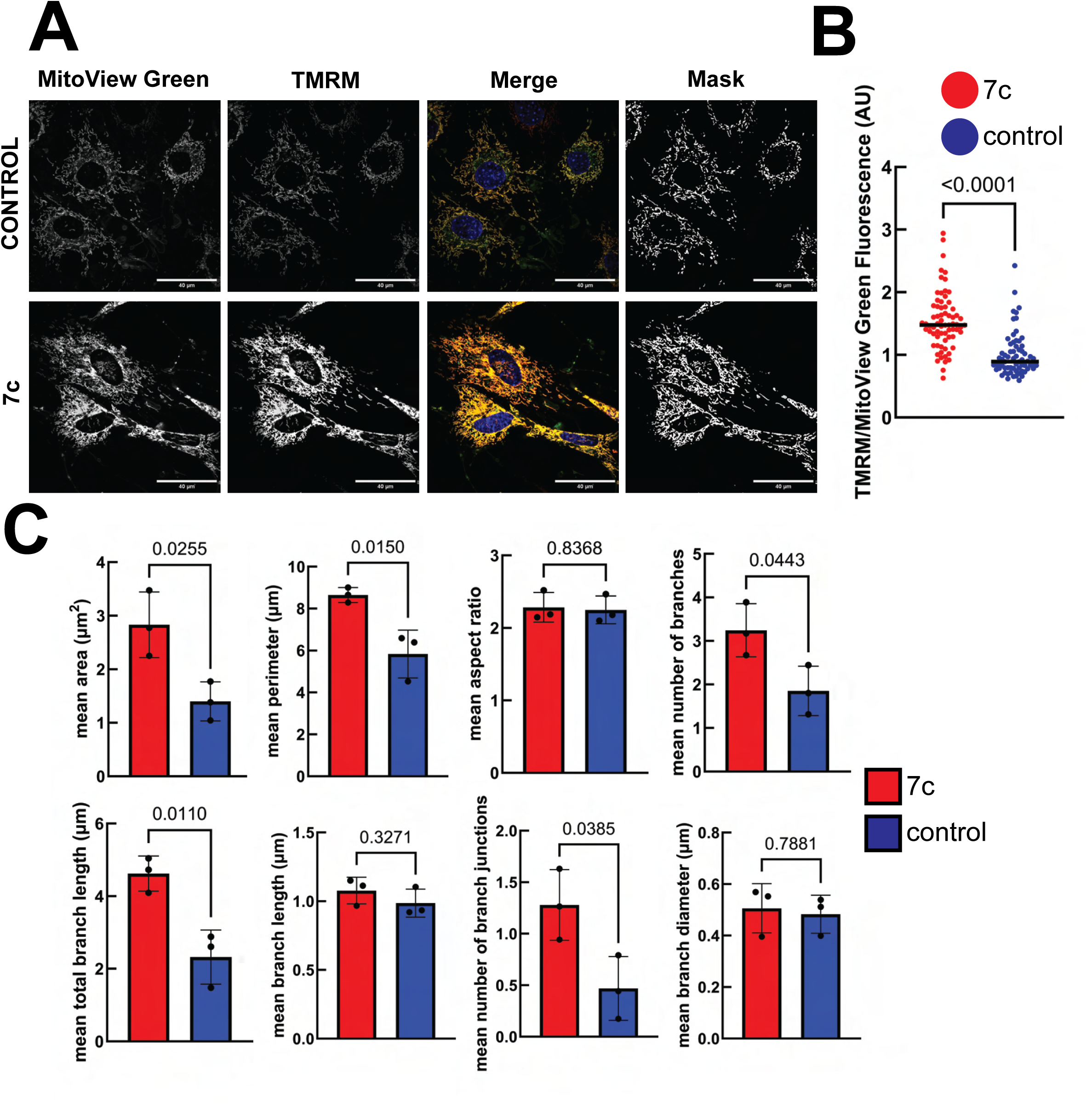
Effect of 7c treatment on mitochondrial 2D morphological networks in mouse fibroblasts. *A. MitoView Green and TMRM staining in live cells.* 2D masks (right panels) were generated by thresholding using the MitoAnalyzer FIJI plug-in (46). Scale bars: 40 µm. *B. Quantification of mitochondrial transmembrane potential.* TMRM signal was normalized to account for changes in mitochondrial mass by dividing by the MitoView Green signal. Bars represent sample means. P-value was determined by two-tailed unpaired t-test. n = 50 images per treatment group from 3 independent biological replicates. *C. Measured mitochondrial 2D parameters.* P-values were determined by two-tailed unpaired t-test. n = 3 independent biological replicates per treatment. Error bars represent sample means ± standard deviation (SD).

**Figure 2:**
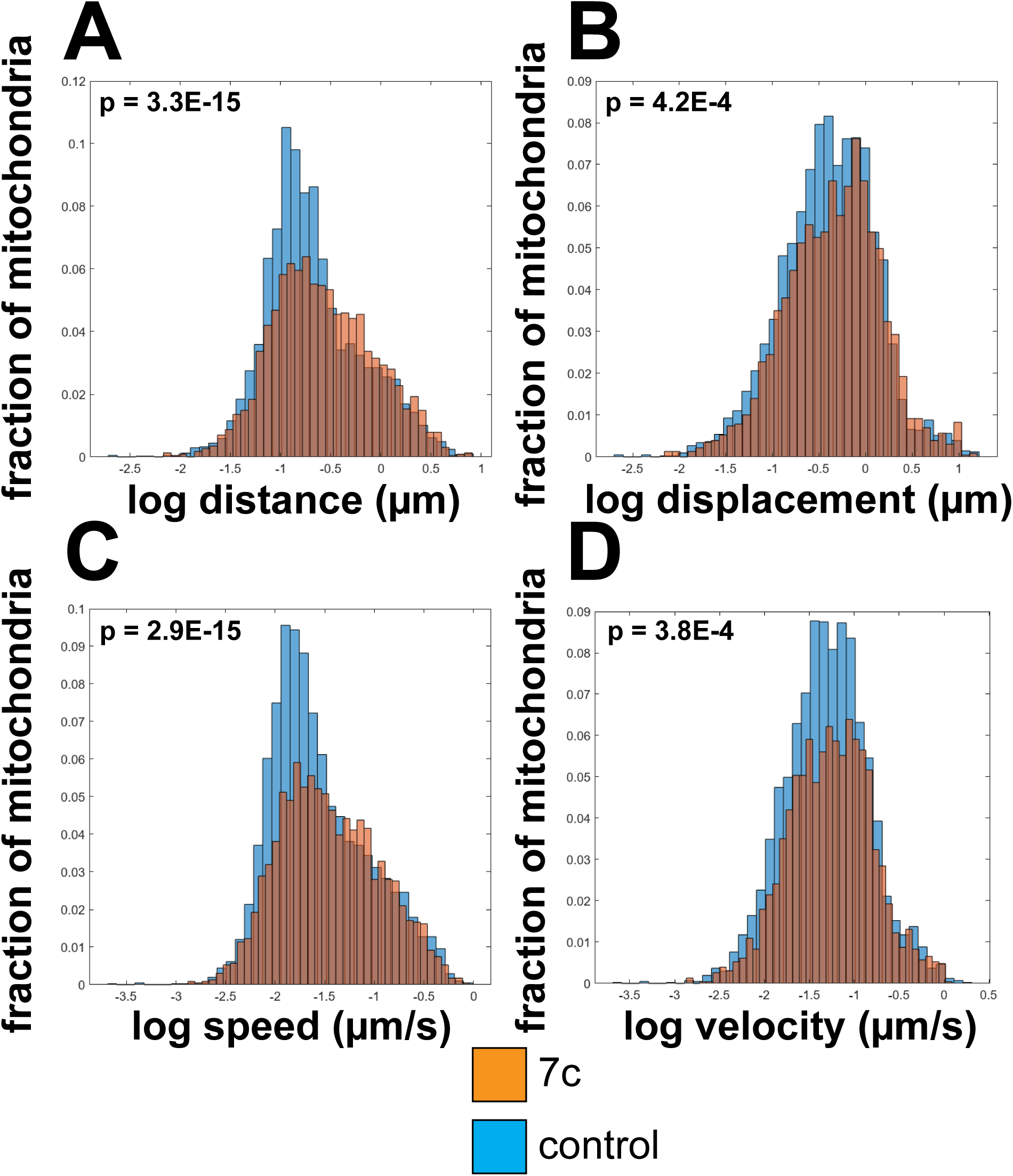
Effect of 7c treatment on mitochondrial 2D dynamics. *A. Distance B. Displacement C. Speed D. Velocity.* P-values were determined by one-way ANOVA using the MitoMeter MATLAB plug-in (47). Data represents n = 15 movies per treatment group across 3 independent biological replicates.

To more precisely measure the effect of 7c partial chemical reprogramming on mitochondrial morphology, we fixed and stained cells for Tom20 and collected 200 nm z-stacks. We then generated 3D masks of the mitochondrial networks using the MitoAnalyzer FIJI plug-in (46) (**Fig. 3A**). Control experiments using pre-treatment with a mitochondrial uncoupler (CCCP, **Supplemental Fig. S1**) demonstrated that dissipation of the mitochondrial transmembrane potential causes a significant decrease in mitochondria size and shape and further disrupts their branching networks. Following 7c treatment, we observed significant increases in mitochondrial volume, surface area, number of branches, and total branch length (**Fig. 3B**). Taken together, these data further verified that partial chemical reprogramming induces significant effects on mitochondrial function, morphology, and dynamics.

**Figure 3:**
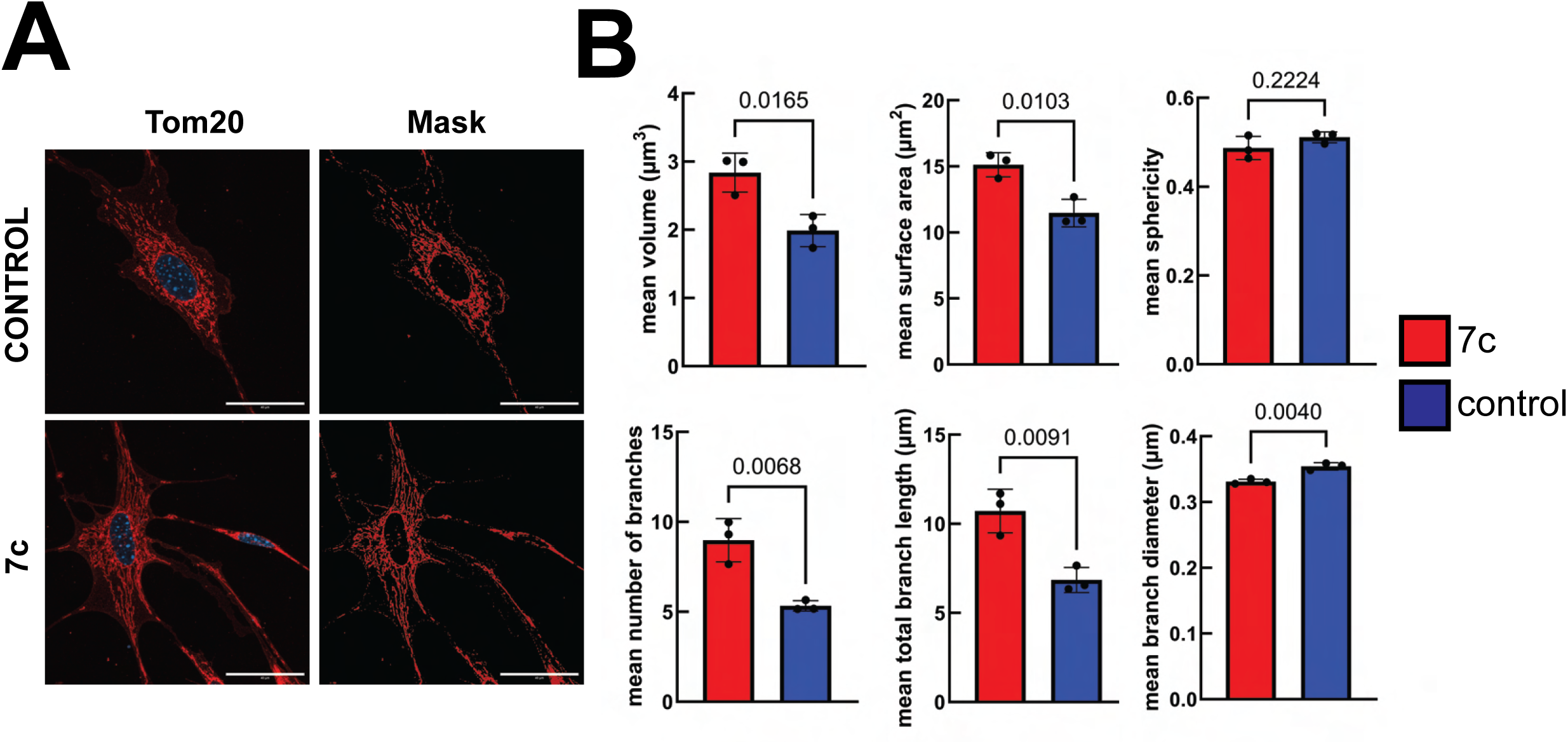
Effect of 7c treatment on mitochondrial 3D morphological networks. *A. 3D projections of Tom20 staining.* 3D projections of Tom20 staining (left panels) and 3D projections of masks (right panels) generated using the MitoAnalyzer FIJI plug-in (46). Scale bars: 40 µm. Z-stacks were obtained using a 0.2 µm step size. *B. Measured 3D mitochondrial parameters.* P-values were determined by two-tailed unpaired t-test. n = 3 independent biological replicates per treatment. Error bars represent sample means ± SD.

As cristae structure and density is closely associated with mitochondrial bioenergetic function (58), we then used transmission electron microscopy (TEM) to ascertain the impact of 7c treatment on cristae morphology. In control cells, we observed typical cristae morphology characterized by tightly stacked, thin lamella that fully cross the mitochondrial minor axes (**Fig. 4A**). However, with 7c-treated cells, we noticed a larger proportion of mitochondria with circular and/or onion-like cristae (red arrows). Although not statistically significant, we also saw a trend towards increased mitochondrial area and perimeter following 7c treatment (**Fig. 4B**). Unexpectedly, we also observed an increase in the number of lipid droplets (purple arrows) following partial chemical reprogramming (**Fig. 4A**). We validated our findings by quantifying oil red o staining in control-, 2c-, and 7c-treated cells (**Fig. 4C**). As expected, we observed significant increases in oil red o staining following 7c treatment compared to control cells, whereas 2c treatment caused an even greater amount of intracellular lipid droplets to form. Given these results, we then re-analyzed our proteomics and phosphoproteomics datasets (43) to look specifically at differences in protein expression following 7c treatment relative to 2c treatment. In both fibroblasts isolated from young (4-month-old) and old (20-month-old) C57BL/6J male mice, we observed a similar higher abundance of OXPHOS proteins, and a lower abundance of cell cycle proteins, after 7c treatment relative to 2c treatment (**Fig. 5A**). At the phosphoproteome level, we noticed key differences in the effects of 2c and 7c on chromatin organization and ion transport, particularly in old mouse fibroblasts (**Fig. 5B**). Based on our previous observations that only 7c significantly increases cellular spare respiratory capacity (43), we then specifically re-analyzed the phosphoproteomic data to assess differential phosphorylation in proteins related to the mitochondrion, lipolysis, structural muscle constituents, and the spliceosome, between 2c and 7c treatments in young and old fibroblasts (**Fig. 5C**). We found that biological processes related to lipolysis and structural muscle constituents are more phosphorylated in 2c-treated fibroblasts compared to 7c-treated fibroblasts, both in young and old cells. Conversely, proteins related to the mitochondrion are significantly more phosphorylated in old 7c-treated fibroblasts compared to old 2c-treated fibroblasts. Therefore, we determined that the 7c and 2c cocktails differ primarily in their impact on lipid droplet formation, mitochondrial OXPHOS, and cell proliferation, which all may be interrelated.

**Figure 4:**
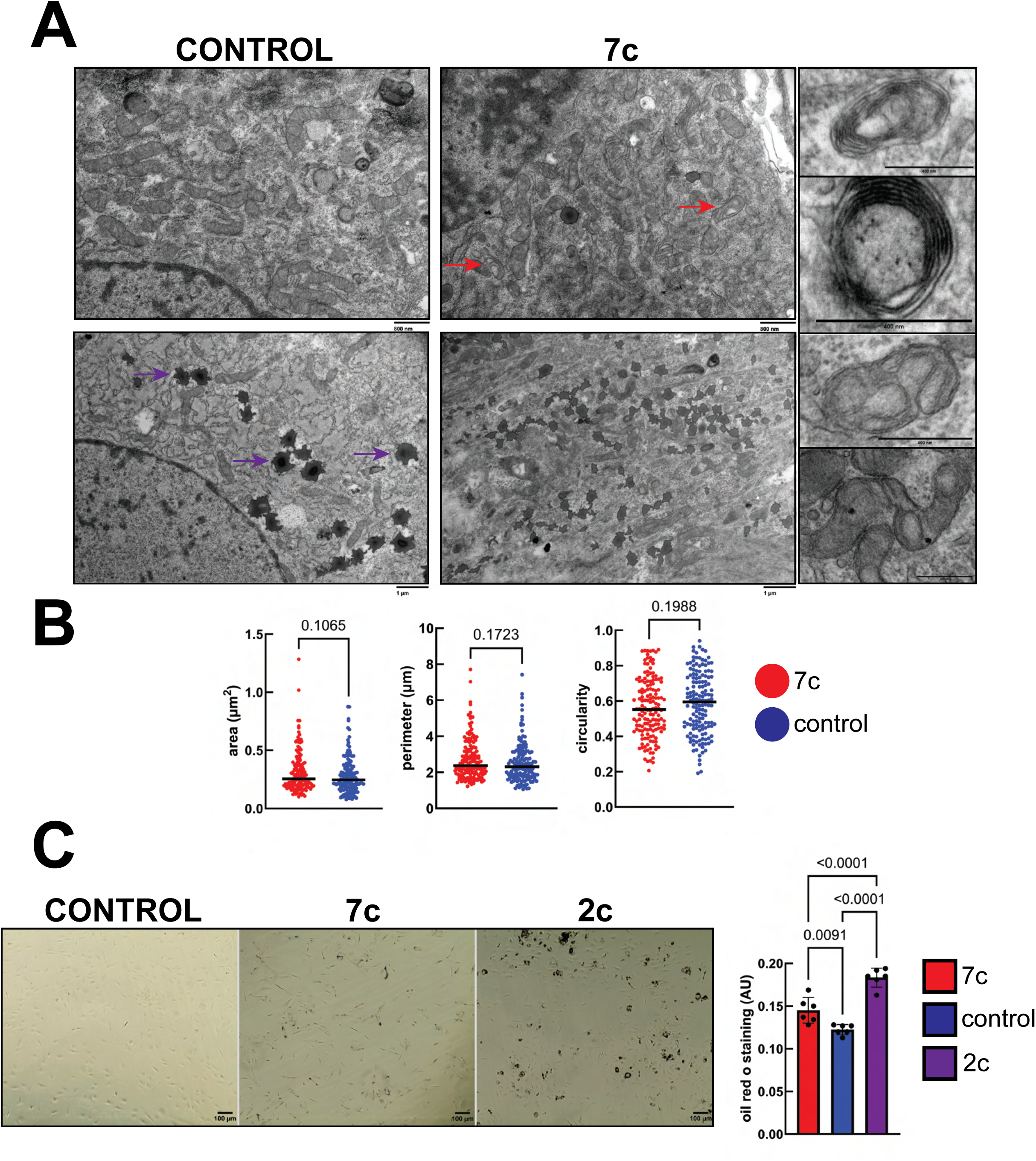
Effect of 7c treatment on mitochondrial cristae morphology and lipid droplet formation. *A. TEM images of lead citrate-stained ultrathin fibroblast cell sections.* Red arrows indicate mitochondria with abnormal/circular cristae morphology. Purple arrows indicate example lipid droplets. Single examples of atypical cristae morphology are shown to the right. *B. Measured mitochondrial parameters.* Bars represent sample means. P-values were determined by two-tailed unpaired t-test. n = 100 randomly selected mitochondria per treatment from images taken of 3 independent biological replicates. *C. Oil red o staining.* Brightfield images of cells stained with oil red o (left panels), and quantification of amount of oil red o staining (right panel). P-values were determined by one-way ANOVA and Tukey’s post-hoc test. n = 6 independent biological replicates. Error bars represent sample means ± SD.

**Figure 5:**
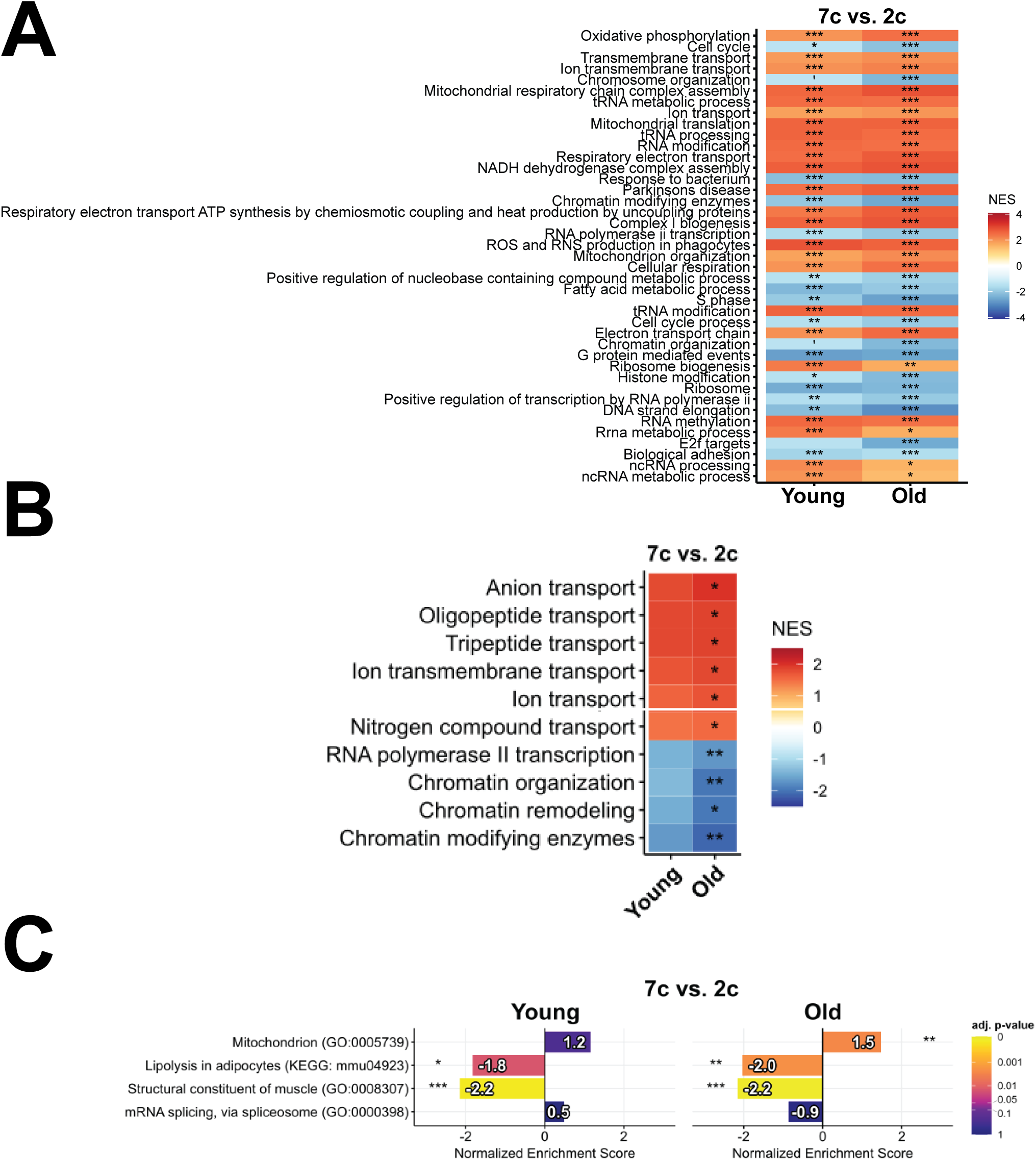
Comparison of 7c vs. 2c treatment on protein and phosphoprotein abundance. *A. Protein enrichment analysis.* Positive and negative normalized enrichment scores (NES) indicate that the biological process is upregulated or downregulated at the protein level in 7c-treated cells compared to 2c-treated cells, respectively. Comparisons were performed using proteomics data obtained from fibroblasts isolated from Young (4-month-old) and Old (20-month-old) male C57BL/6J mice (43). We show the 25 most significant protein sets in Young and Old. P-values were adjusted for multiple comparisons using the Benjamini-Hochberg false discovery rate (FDR) method. n = 3 biological replicates per treatment group. ‘adjusted p-value < 0.1, *adjusted p-value < 0.05, **adjusted p-value < 0.01, ***adjusted p-value < 0.001. *B. Phosphoprotein enrichment analysis.* Positive and negative normalized enrichment scores (NES) indicate that the proteins in the biological process are more phosphorylated or dephosphorylated in 7c-treated cells compared to 2c-treated cells, respectively. Comparisons were performed using phosphoproteomics data obtained from fibroblasts isolated from Young (4-month-old) and Old (20-month-old) male C57BL/6J mice (43). We show the 10 significant protein sets in old fibroblasts; no protein sets are significant in young fibroblasts at a 5% FDR significance threshold. P-values were adjusted for multiple comparisons using the Benjamini-Hochberg FDR method. n = 3 biological replicates per treatment group. *adjusted p-value < 0.05, **adjusted p-value < 0.01. *C. Targeted phosphoprotein enrichment analysis.* Positive and negative normalized enrichment scores (NES) indicate that the proteins in the four pre-selected biological process are more phosphorylated or dephosphorylated in 7c-treated cells compared to 2c-treated cells, respectively. Numbers depicted in the bars represent the NES. P-values were adjusted for multiple comparisons using the Benjamini-Hochberg method. n = 3 biological replicates per treatment group. *adjusted p-value < 0.05, **adjusted p-value < 0.01, ***adjusted p-value < 0.001.

We then sought to investigate whether 7c partial chemical reprogramming could affect molecular biomarkers of aging in a genetically heterogeneous mouse model. We chose to use the 7c cocktail because we previously showed that it robustly decreased the transcriptomic and epigenetic age of mouse fibroblasts (43). To address the challenge of the 7c compounds’ high insolubility in water and the potential toxicity of larger amounts of DMSO administered *in vivo*, we developed a system for stable, long-term delivery of the 7c cocktail over a 1-month period using subcutaneously (SQ) implantable osmotic minipumps (**Fig. 6A**). A solvent of 50% DMSO and 50% PEG_300_ effectively solubilized the 7c cocktail and was compatible with the pumps. Additionally, the volume of pure DMSO (which is toxic (59, 60)) delivered per day was less than µL, and the minor surgical procedure is well-tolerated by aged mice (50, 61). We delivered 0.1 mg/kg/day of 6c (repsox, tranylcypromine, DZNep, forskolin, CHIR99021, TTNPB) and 1 mg/kg/day of valproate to 12-month-old male UM-HET3 mice for a period of 28 days (**Table S1**), with control mice receiving pumps filled only with solvent. After 28 days of treatment, the mice were euthanized, and RNA extracted from their livers and kidneys were analyzed using bulk mRNA-seq.

**Figure 6:**
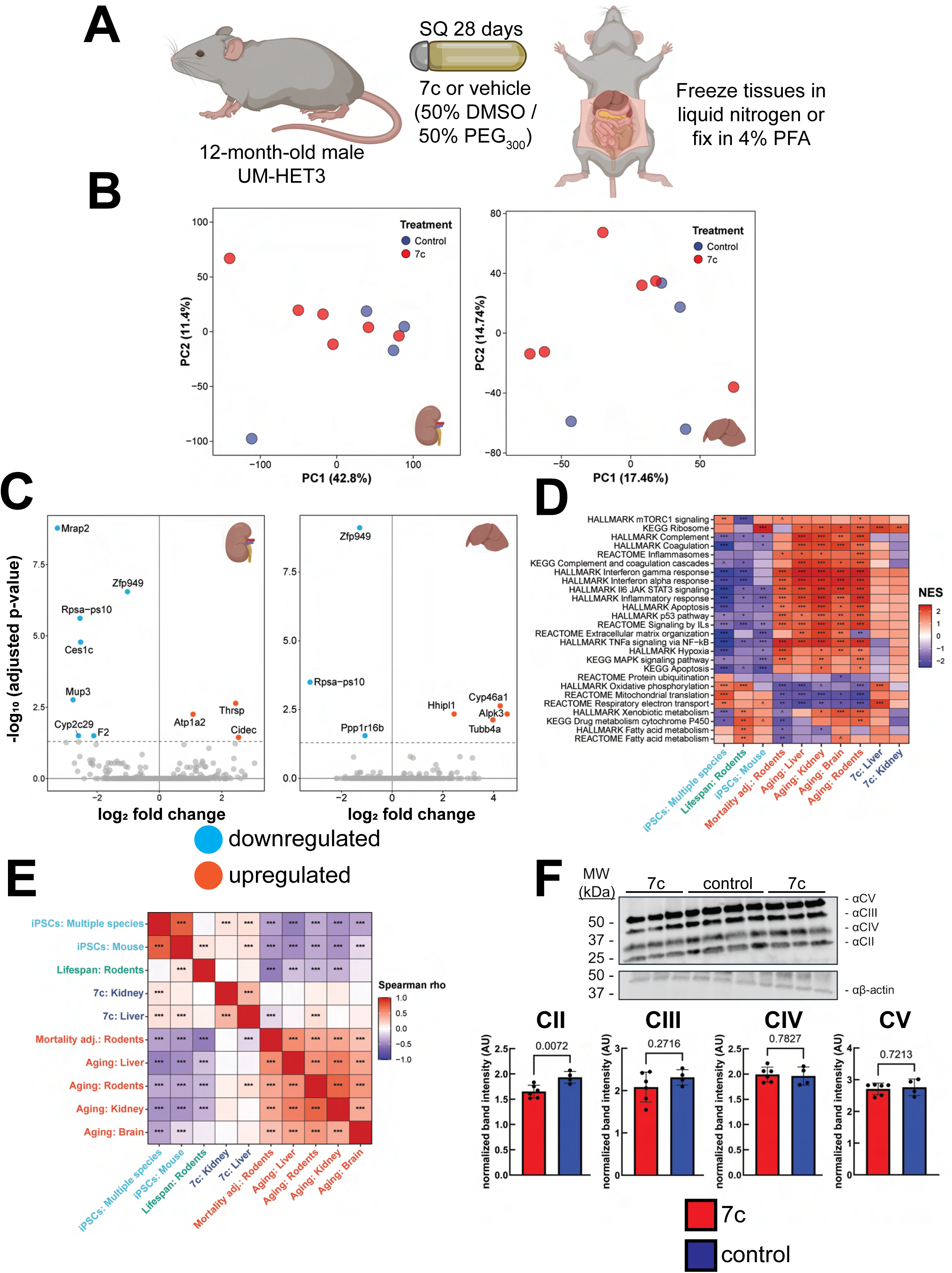
Effect of 28 days low-dose 7c treatment on gene and protein expression in male UM-HET3 mice. *A. Experimental strategy for in vivo partial chemical reprogramming. B. Principal component analysis of bulk mRNA-seq data.* Shown here for mRNA-seq data obtained from kidney (left) and liver (right) samples. n = 4-6 mice per treatment. *C. Detected differentially expressed genes following 7c treatment.* Significantly (adjusted p-value < 0.05) downregulated (cyan) and upregulated (red) genes are labeled for kidney (left panel) and liver (right panel) tissues. *D. Gene set enrichment analysis (GSEA) of pathways affected by 7c treatment (blue).* Shown here compared to signatures of aging and mortality (red), iPSCs (cyan), and expected maximum lifespan (green). Gene sets from KEGG, REACTOME, and HALLMARKS ontologies were utilized. The complete list of significantly affected functions is in Tables S4-S5. NES: normalized enrichment score. ^adjusted p-value < 0.1, *adjusted p-value < 0.05, **adjusted p-value < 0.01, ***adjusted p-value < 0.001. *E. Spearman correlation analysis of enriched pathways perturbed by 7c treatment (blue).* Shown here compared to signatures of aging and mortality (red), iPSCs (cyan), and expected maximum lifespan (green). Pairwise Spearman correlations were calculated based on NES values determined for each signature via GSEA. ***adjusted p-value < 0.001. *F. Effect of 7c treatment on abundance of OXPHOS proteins.* Bands for OXPHOS proteins were ormalized to their respective β-actin bands. P-values were determined by two-tailed unpaired t-test. n = 4-6 mice per treatment.

To evaluate the *in vivo* efficacy of 7c, we first utilized transcriptomic biomarkers of aging to analyze the liver and kidney bulk mRNA-seq data. By principal component analysis (**Fig. 6B**) of kidney (left panel) and liver (right panel) mRNA-seq samples, we did not observe separation between 7c- and vehicle-treated animals on either principal component 1 or 2. After performing a differential expression analysis (**Tables S2-S3**), we observed very few differentially expressed genes in both the livers and kidneys with only a downregulation of *Zfp949* and *Rpsa-ps10* being shared between the two tissues (**Fig. 6C**). We then performed gene set enrichment analysis (GSEA) to determine if 7c treatment impacted the expression of gene sets associated with mammalian aging, lifespan regulation, and OSKM reprogramming (**Tables S4-S5, Fig. 6D**). The transcriptional changes induced by 7c in both the kidney and liver (blue) did not globally resemble the functional signatures of aging (red), iPSCs (cyan), or established lifespan-extending interventions (green). At the level of individual pathways, the only consistently and significantly upregulated gene sets after 28 days of 7c treatment were those related to OXPHOS and ribosomal organization. We further performed a Spearman correlation analysis comparing the gene expression changes induced by 7c with reference signatures at the level of enriched functions (**Fig. 6E**). Transcriptional changes induced by 7c in the liver exhibited modest positive correlation with iPSCs and rodent aging signatures, and a negative correlation with signatures of rodent mortality. Furthermore, we were unable to detect an impact of 7c treatment on kidney or liver transcriptomic age using either rodent chronological or mortality transcriptomic clocks (**Supplemental Fig. S2**). Although OXPHOS genes were positively enriched at the transcriptomic level in the liver, we did not observe a corresponding increase in OXPHOS protein abundance after 28 days with 7c treatment (**Fig. 6F**). Thus, we concluded that at this first dosage tested, 7c treatment is safely administered but does not lead to robust changes in mitochondrial OXPHOS or transcriptomic biomarkers of biological age.

Since we didn’t observe any evidence of lowered transcriptomic age or upregulation of mitochondrial OXPHOS with this first treatment regiment, we decided to increase the dosage to 0.5 mg/kg/day for 6c and 50 mg/kg/day for valproate (**Fig. 7A**). However, this dose resulted in a rapid decrease in body weight and body condition score in ∼5-6 days that required euthanasia. A follow-up trial using an intermediate dosage of 0.25 mg/kg/day for 6c and 10 mg/kg/day for valproate similarly resulted in a significant decrease in body weight after 7 days (**Fig. 7B**). Gross necropsy did not reveal any readily apparent features (e.g. tumors, hemorrhaging, liver failure) that may have contributed to the drug-induced toxicity. Therefore, we then sectioned the fixed liver and kidney tissues and stained both with hematoxylin and eosin, Masson’s trichrome, and oil red o. In the livers (**Fig. 7C**) of animals treated with 7c, we did not observe increased collagen staining indicative of fibrosis, or any apoptotic bodies being formed (33). However, we did notice a strong increase in oil red o staining that was further validated by quantification of the stained area. Similarly, we did not observe an increase in fibrosis in the kidneys of the 7c-treated animals (**Supplemental Fig. S3**) but also saw a strong increase in overall oil red o staining (**Fig. 7D**). In the animals treated with vehicle, we observed age-related glomerular changes and positive lipid staining (black arrows). However, for the animals treated with 7c, we noticed a significant decrease of oil red o staining in the glomeruli. Therefore, we concluded that 7c treatment can cause the accumulation of triglycerides in multiple organs and alter kidney microscopic anatomy.

**Figure 7:**
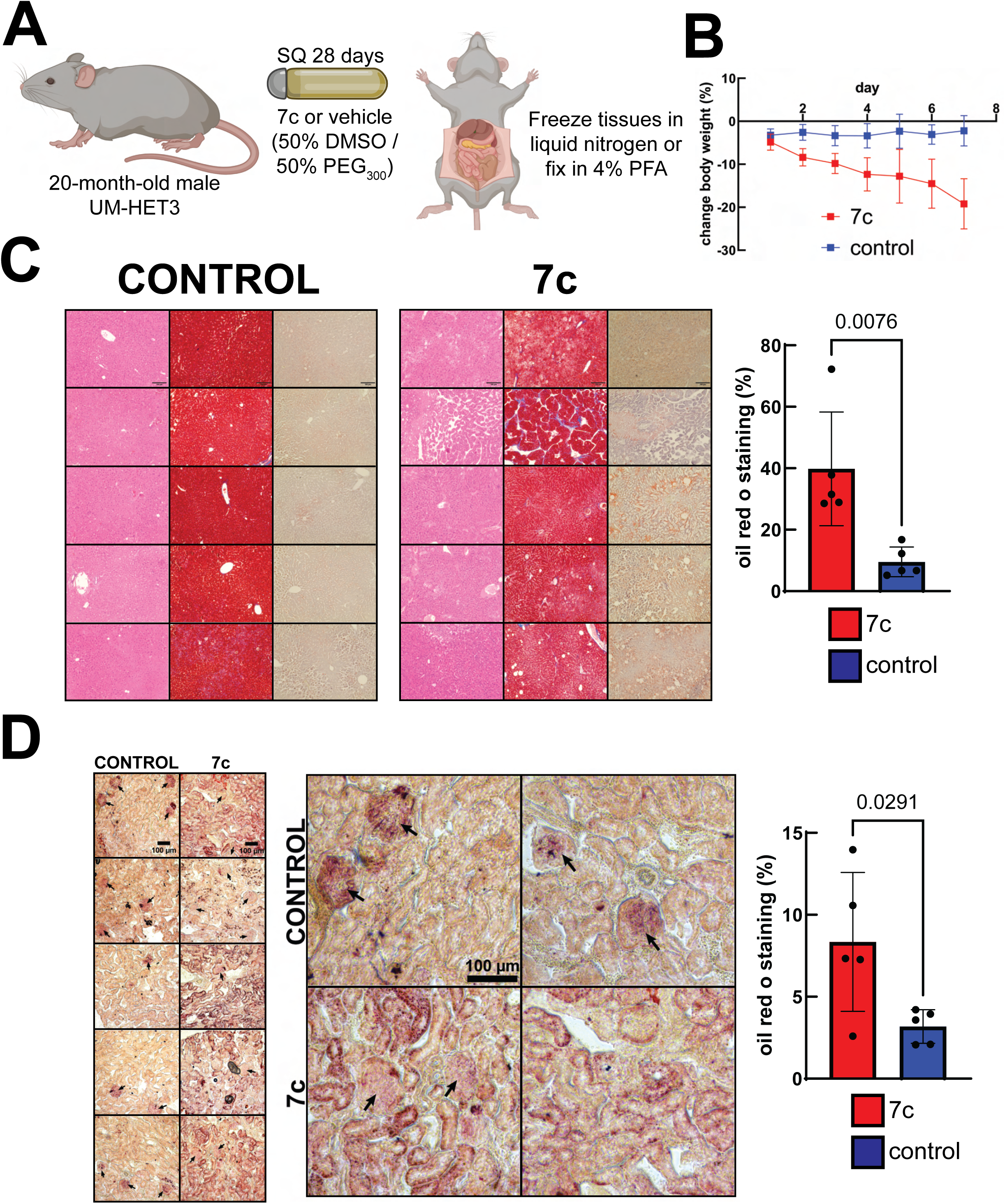
Effect of high-dose 7c treatment in male UM-HET3 mice. *A. Experimental strategy for in vivo partial chemical reprogramming. B. Effect of high-dose 7c treatment on body weight.* Data points represent sample means ± SD. n = 6 mice per treatment. *C. Effect of high-dose 7c treatment on liver histology.* Liver sections were stained with hematoxylin and eosin (left panels), Masson’s trichrome (center panels), and oil red o (right panels). Each row represents images obtained from stained tissues of a single animal. Scale bars: 100 µm. n = 5 mice per treatment. Error bars represent sample means ± SD. P-value was determined by two-tailed unpaired t-test. *D. Effect of high-dose 7c treatment on kidney histology.* Kidney sections were stained with oil red o. Each panel represents an image obtained from stained tissue of a single animal. Scale bars: 100 µm. Arrows indicate glomeruli. n = 5 mice per treatment. P-value was determined by two-tailed unpaired t-test.

Due to the changes we observed in glomeruli lipid staining following 7c treatment, we then evaluated 7c treatment for its impact on markers of acute renal injury. We first stained kidney sections for LTL, which is a general marker for proximal tubule brush border integrity (62), and KIM-1 (**Fig. 8A**), which is upregulated in the proximal tubule after injury/ischemia (63). In kidneys from control animals, very few proximal tubules were positive for KIM-1 expression. Although two 7c-treated animals showed pronounced increases in KIM-1-stained proximal tubules, the effect across the entire treatment group was not significant. However, when we stained with phalloidin, we noticed a significant loss in polarized tubules with apical F-actin in the 7c-treated animals (**Fig. 8B**). It has been previously reported that the cytoskeleton of proximal tubules is disrupted after ischemia (64). Finally, to assess overall kidney function, we measured serum creatinine levels using mass spectrometry (**Fig. 8C**). Creatinine levels were generally low across both treatment groups (65); however, serum creatinine levels in the 7c-treated animals were slightly elevated with marginal statistical significance. Therefore, we concluded that high-dose 7c treatment can cause acute injury reminiscent of ischemia.

**Figure 8:**
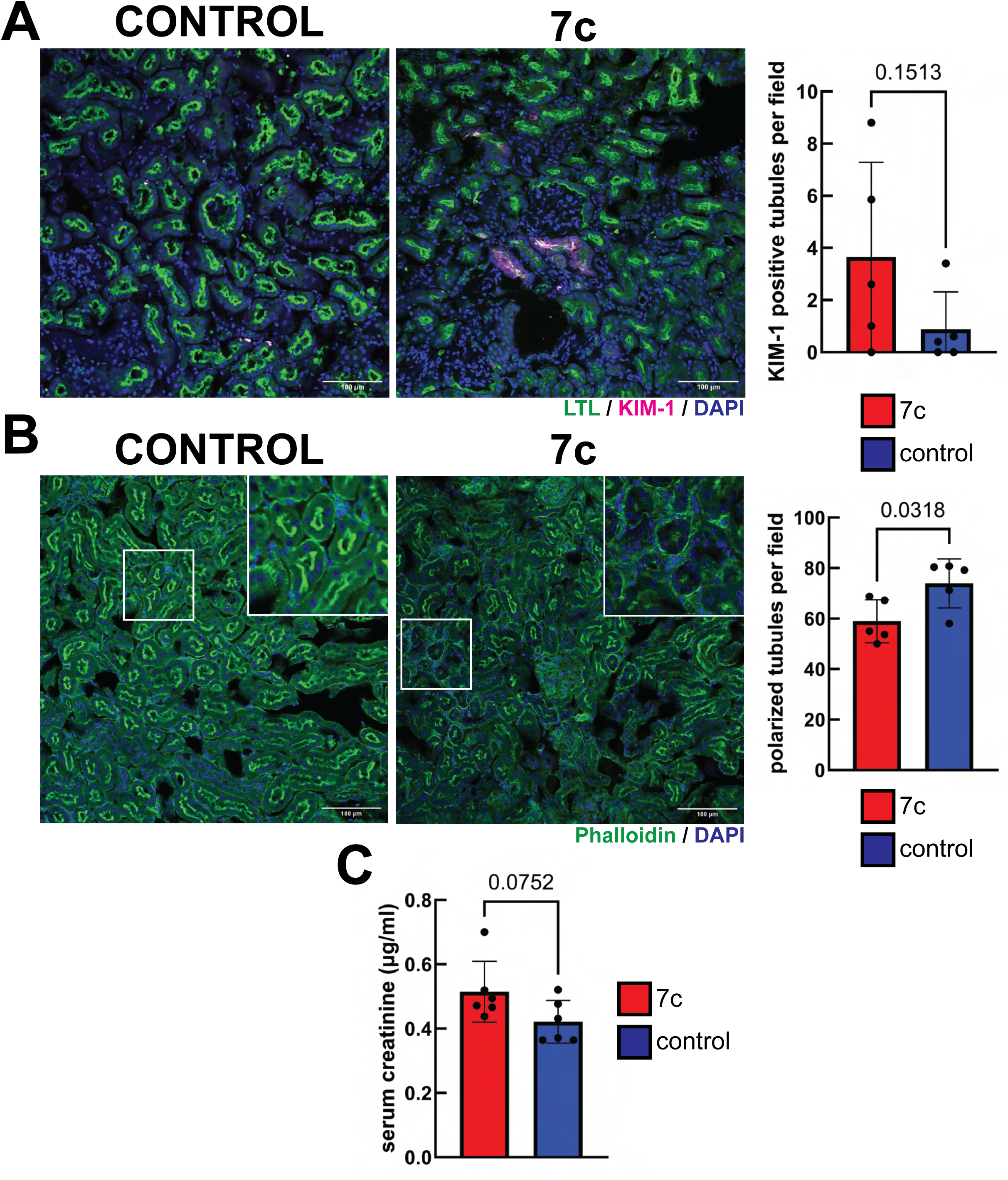
Effect of high-dose 7c treatment on markers of acute kidney injury. *A. Effect on renal tubular injury.* Representative images of KIM-1-and LTL-stained kidney sections from control and 7c-treated mice. 5-7 images were acquired per mouse. P-values were determined by two-tailed unpaired t-test. n = 5 mice per treatment. Error bars represent sample means ± SD. Scale bars: 100 µm. *B. Effect on renal tubular polarization.* Representative images of phalloidin-stained kidney sections from control and 7c-treated mice. 5-9 images were acquired per mouse. P-values were determined by two-tailed unpaired t-test. n = 5 mice per treatment. Error bars represent sample means ± SD. Scale bars: 100 µm. *C. Effect on serum creatinine levels.* P-value was determined by two-tailed unpaired t-test. n = 6 mice per treatment. Error bars represent sample means ± SD.

## DISCUSSION

We assessed the impact of 7c partial chemical reprogramming on mitochondrial morphology and dynamics in mouse fibroblasts and measured the impact of 7c treatment on organismal biological age and health using genetically diverse UM-HET3 male mice. Consistent with previous reports, we have determined that 7c partial chemical reprogramming causes significant changes to mitochondrial interaction networks, movement dynamics, and morphology. In tandem, we observed that treatment with both 7c and 2c causes a significant increase in lipid droplet formation *in vitro,* with 2c treatment having an even more pronounced effect compared to 7c treatment in fibroblasts. This result was further supported by our previous phosphoproteomics data, which demonstrated that relative to 7c treatment, 2c treatment significantly dephosphorylates proteins involved in lipolysis. When we treated male UM-HET3 mice with a low dose of the 7c cocktail for 28 days, we did not observe significant effects on kidney or liver transcriptomic age, overall gene expression, or on the expression of mitochondrial OXPHOS proteins. Higher doses of 7c, in contrast, led to a rapid loss of body weight and lowered body condition scores that required euthanasia. Therefore, partial chemical reprogramming currently seems unable to induce more youthful gene expression profiles in mammalian tissues without causing toxicity.

While Masson’s trichrome and H&E staining did not reveal the presence of fibrosis or apoptotic bodies, we did observe significant lipid accumulation in the livers (similar to early-stage steatosis (66)) and kidneys of 7c-treated animals. We also noticed a slight elevation in biomarkers of acute kidney injury, including cytoskeletal changes in proximal tubules and some 7c-treated animals with increased proximal tubule KIM-1 expression. Thus, our current research suggests that this rapid and severe accumulation of intracellular lipids during partial chemical reprogramming may contribute to the signs of toxicity we observed in our mouse models.

It is important to note the strong differences between reprogramming and chemical reprogramming. On one hand, gene expression signatures of OSKM reprogramming reveal a suppression of mesenchymal genes, increased proliferation, and a metabolic switch to glycolysis during the early phases of reprogramming (67). Moreover, there is decreased and increased expression of genes related to inflammation and DNA repair, respectively (53). Despite these transcriptional changes, the epigenome and histone trimethylation levels of somatic cells are largely maintained until later in the reprogramming process when the expression of epithelial genes is activated (67). On the other hand, chemical reprogramming is marked by global demethylation along with an early transition to a plastic extraembryonic endoderm (XEN)-like cells during stage I (68). This occurs in tandem with upregulated OXPHOS, increased reactive oxygen species production, and reduced cell proliferation (42, 43). Notably, the expression of most pluripotency markers is not activated until stage II, in which *Oct4* activation occurs simultaneously with a gradual downregulation of XEN genes (43, 68). Therefore, these data suggest that partial reprogramming and partial chemical reprogramming rejuvenate cells through entirely disparate mechanisms and as a result, may present distinct advantages and challenges for their applications *in vivo*.

Significant evidence exists that supports the interaction of mitochondria with lipid droplets. These interactions may be crucial for regulating fatty acid metabolism and overall energy homeostasis under both basal and stress conditions (69-71). Thus, in the case of partial chemical reprogramming with 7c, it may be that the strong increase in lipid droplet formation is necessary to mediate the increase in mitochondrial bioenergetic capacity, and vice versa. With 2c, lipid accumulation in contrast is higher, which may make this cocktail even more toxic *in vivo*, yet it does not lead to robust changes in mitochondrial OXPHOS function. While current data cannot causally relate the increase in mitochondrial OXPHOS following 7c partial chemical reprogramming to the reduction in cellular transcriptomic and epigenetic age, we do note that the upregulation of mitochondrial/OXPHOS genes contributes significantly to the observed decrease in transcriptomic age induced by this cocktail in cells (**Supplemental Fig. S4**). Given that the lipid droplets being formed during partial chemical reprogramming appear to cause toxicity, one might assume that modification and/or removal of compound(s) primarily responsible for this effect could lead to a safer cocktail that can also lower biological age *in vivo*. However, the possibility cannot be ignored that the lowered cellular biological age in response to 7c treatment may be dependent on this increase in lipid droplet formation. Therefore, further experimentation is required to determine whether or not the rejuvenating and toxic effects of partial chemical reprogramming can be separated.

## LIMITATIONS AND FUTURE DIRECTIONS

The *in vivo* studies described herein were limited in the number of biological replicates and to only one sex. Therefore, it is unknown if 7c treatment would produce similar effects in female mice. For future studies, it would be important to determine which compound(s) are primarily responsible for the observed toxic effects, and to see if a cocktail without these compound(s) could still have pronounced effects on cell biological age.

## ACKNOWLEDGEMENTS

The authors would like to thank the Microscopy Resources on the North Quad (MicRoN) core at Harvard Medical School (HMS) for providing access and training to the confocal microscopes used in this study, the HMS Electron Microscopy Facility for preparing the EM grids and providing access to the TecnaiG^2^ Spirit BioTWIN electron microscope, and the Dana-Farber/Harvard Cancer Center Rodent Histopathology Core for performing some of the tissue embedding, sectioning, and staining.

## DATA AVAILABILITY

mRNA-seq data will be made publicly available no later than the date of initial publication. All other supporting data is available as supplemental data files and/or upon request from the corresponding author.

## CONTRIBUTIONS

W.M., V.N.G. conceptualization; W.M., C.M., A.T., Y.U., L.J.E.G., T.I., E.N., J.V.B., V.N.G. methodology; W.M., C.M., Y.U. data curation; W.M., C.M., A.T., Y.U., L.J.E.G., T.I. formal analysis; W.M., C.M., A.T., Y.U., L.J.E.G., T.I., E.N. validation; W.M., C.M., A.T., Y.U., L.J.E.G. visualization; W.M., writing – original draft; W.M., C.M., A.T., Y.U., L.J.E.G., E.N., T.I., J.V.B., V.N.G. writing – review and editing; J.V.B., V.N.G. supervision; J.V.B., V.N.G. project administration; V.N.G. funding acquisition.

**Supplemental Figure S1: Effect of uncoupler treatment on mitochondrial morphology and networks**

Mouse fibroblasts were treated with 25 µM carbonyl cyanide m-chlorophenylhydrazone (CCCP) for 30 minutes prior to fixation, permeabilization, and staining for Tom20. Mitochondrial parameters were measured from 3D z-stacks (0.5 µm step size) using the MitoAnalyzer FIJI plug-in (46). P-values were determined by two-tailed unpaired t-test. Error bars represent sample means ± SD.

**Supplemental Figure S2: Effect of *in vivo* partial chemical reprogramming on liver and kidney tissue transcriptomic age**

Chronological (upper panels) and mortality (lower panels) rodent multi-tissue transcriptomic clocks based on Bayesian Ridge model were used to predict the transcriptomic age of kidney (left panels) and liver (right panels) tissues from 7c- and control-treated animals using the bulk mRNA-seq data presented in Fig. 6. n = 4-6 mice per treatment.

**Supplemental Figure S3: Additional kidney histology images**

5 µm sections of paraformaldehyde fixed, paraffin embedded kidney tissue from either 7c- or control-treated animals were stained with Masson’s trichrome (left panels) and hematoxylin and eosin (right panels). Each row contains images from an individual mouse. Scale bars: 100 µm. n = 4 mice per treatment.

**Supplemental Figure S4: Modular transcriptomic age of fibroblasts treated with 2c and 7c** Effect of 2c and 7c treatment on cell transcriptomic age predicted by biological module-specific multi-tissue clocks (19). The effect of 7c and 2c on cell gene expression was measured relative to control cells treated with vehicle. mRNA-seq data was obtained from (43). ^adjusted p-value < 0.1, *adjusted p-value < 0.05, **adjusted p-value < 0.01, ***adjusted p-value < 0.001. n = 8 biological replicates per treatment group.

## REFERENCES

1. López-Otín, C., Blasco, M. A., Partridge, L., Serrano, M., and Kroemer, G. (2013) The Hallmarks of Aging Cell 153, 1194-1217 10.1016/j.cell.2013.05.039

2. Ferrucci, L., and Kuchel, G. A. (2021) Heterogeneity of Aging: Individual Risk Factors, Mechanisms, Patient Priorities, and Outcomes Journal of the American Geriatrics Society 69, 610–612 10.1111/jgs.17011

3. Bou Sleiman, M., Roy, S., Gao, A. W., Sadler, M. C., von Alvensleben, G. V. G., Li, H., et al. (2022) Sex- and age-dependent genetics of longevity in a heterogeneous mouse population Science 377, eabo3191 doi:10.1126/science.abo3191

4. Yang, M., Harrison, B. R., and Promislow, D. E. L. (2023) Cellular age explains variation in age-related cell-to-cell transcriptome variability Genome Res 33, 1906-1916 10.1101/gr.278144.123

5. Shadel, G. S., Adams, P. D., Berggren, W. T., Diedrich, J. K., Diffenderfer, K. E., Gage, F. H. et al. (2021) The San Diego Nathan Shock Center: tackling the heterogeneity of aging Geroscience 43, 2139-2148 10.1007/s11357-021-00426-x

6. Hägg, S., and Jylhävä, J. (2021) Sex differences in biological aging with a focus on human studies eLife 10, e63425 10.7554/eLife.63425

7. Lu, Y. R., Tian, X., and Sinclair, D. A. (2023) The Information Theory of Aging Nat Aging 3, 1486-1499 10.1038/s43587-023-00527-6

8. Gladyshev, V. N. (2014) The free radical theory of aging is dead. Long live the damage theory! Antioxid Redox Signal 20, 727–731 10.1089/ars.2013.5228

9. Gladyshev, V. N., Kritchevsky, S. B., Clarke, S. G., Cuervo, A. M., Fiehn, O., de Magalhães, J. P. et al. (2021) Molecular damage in aging Nature Aging 1, 1096-1106 10.1038/s43587-021-00150-3

10. Gladyshev, V. N. (2016) Aging: progressive decline in fitness due to the rising deleteriome adjusted by genetic, environmental, and stochastic processes Aging Cell 15, 594–602 10.1111/acel.12480

11. Austad, S. N., and Hoffman, J. M. (2018) Is antagonistic pleiotropy ubiquitous in aging biology? Evolution, Medicine, and Public Health 2018, 287–294 10.1093/emph/eoy033

12. Gems, D. (2022) The hyperfunction theory: An emerging paradigm for the biology of aging Ageing Res Rev 74, 101557 10.1016/j.arr.2021.101557

13. Blagosklonny, M. V. (2012) Answering the ultimate question “what is the proximal cause of aging?” Aging (Albany NY) 4, 861–877 10.18632/aging.100525

14. Sierra, F. (2016) The Emergence of Geroscience as an Interdisciplinary Approach to the Enhancement of Health Span and Life Span Cold Spring Harb Perspect Med 6, a025163 10.1101/cshperspect.a025163

15. Li, Z., Zhang, W., Duan, Y., Niu, Y., Chen, Y., Liu, X., et al. (2023) Progress in biological age research Front Public Health 11, 1074274 10.3389/fpubh.2023.1074274

16. Zurbuchen, R., von Däniken, A., Janka, H., von Wolff, M., and Stute, P. (2025) Methods for the assessment of biological age -A systematic review Maturitas 195, 108215 10.1016/j.maturitas.2025.108215

17. Rutledge, J., Oh, H., and Wyss-Coray, T. (2022) Measuring biological age using omics data Nature Reviews Genetics 23, 715–727 10.1038/s41576-022-00511-7

18. Tyshkovskiy, A., Bozaykut, P., Borodinova, A. A., Gerashchenko, M. V., Ables, G. P., Garratt, M. et al. (2019) Identification and Application of Gene Expression Signatures Associated with Lifespan Extension Cell Metab 30, 573–593.e578 10.1016/j.cmet.2019.06.018

19. Tyshkovskiy, A., Kholdina, D., Ying, K., Davitadze, M., Molière, A., Tongu, Y., et al. (2024) Transcriptomic Hallmarks of Mortality Reveal Universal and Specific Mechanisms of Aging, Chronic Disease, and Rejuvenation bioRxiv 2024.2007.2004.601982 10.1101/2024.07.04.601982

20. Lu, A. T., Quach, A., Wilson, J. G., Reiner, A. P., Aviv, A., Raj, K. et al. (2019) DNA methylation GrimAge strongly predicts lifespan and healthspan Aging (Albany NY) 11, 303–327 10.18632/aging.101684

21. Belsky, D. W., Caspi, A., Corcoran, D. L., Sugden, K., Poulton, R., Arseneault, L., et al. (2022) DunedinPACE, a DNA methylation biomarker of the pace of aging eLife 11, e73420 10.7554/eLife.73420

22. Lu, A. T., Fei, Z., Haghani, A., Robeck, T. R., Zoller, J. A., Li, C. Z. et al. (2023) Universal DNA methylation age across mammalian tissues Nature Aging 3, 1144-1166 10.1038/s43587-023-00462-6

23. Ying, K., Liu, H., Tarkhov, A. E., Sadler, M. C., Lu, A. T., Moqri, M. et al. (2024) Causality-enriched epigenetic age uncouples damage and adaptation Nat Aging 4, 231–246 10.1038/s43587-023-00557-0

24. Goeminne, L. J. E., Vladimirova, A., Eames, A., Tyshkovskiy, A., Argentieri, M. A., Ying, K. et al. (2025) Plasma protein-based organ-specific aging and mortality models unveil diseases as accelerated aging of organismal systems Cell Metab 37, 205–222.e206 10.1016/j.cmet.2024.10.005

25. Oh, H. S.-H., Rutledge, J., Nachun, D., Pálovics, R., Abiose, O., Moran-Losada, P. et al. (2023) Organ aging signatures in the plasma proteome track health and disease Nature 624, 164–172 10.1038/s41586-023-06802-1

26. McCrory, C., Fiorito, G., Hernandez, B., Polidoro, S., O’Halloran, A. M., Hever, A. et al. (2020) GrimAge Outperforms Other Epigenetic Clocks in the Prediction of Age-Related Clinical Phenotypes and All-Cause Mortality The Journals of Gerontology: Series A 76, 741–749 10.1093/gerona/glaa286

27. Takahashi, K., and Yamanaka, S. (2006) Induction of pluripotent stem cells from mouse embryonic and adult fibroblast cultures by defined factors Cell 126, 663–676 10.1016/j.cell.2006.07.024

28. Singh, P. B., and Zhakupova, A. (2022) Age reprogramming: cell rejuvenation by partial reprogramming Development 149, 10.1242/dev.200755

29. Paine, P. T., Nguyen, A., and Ocampo, A. (2024) Partial cellular reprogramming: A deep dive into an emerging rejuvenation technology Aging Cell 23, e14039 10.1111/acel.14039

30. Ocampo, A., Reddy, P., Martinez-Redondo, P., Platero-Luengo, A., Hatanaka, F., Hishida, T. et al. (2016) In Vivo Amelioration of Age-Associated Hallmarks by Partial Reprogramming Cell 167, 1719-1733.e1712 10.1016/j.cell.2016.11.052

31. Browder, K. C., Reddy, P., Yamamoto, M., Haghani, A., Guillen, I. G., Sahu, S. et al. (2022) In vivo partial reprogramming alters age-associated molecular changes during physiological aging in mice Nature Aging 2, 243–253 10.1038/s43587-022-00183-2

32. Sahu, S. K., Reddy, P., Lu, J., Shao, Y., Wang, C., Tsuji, M., et al. (2024) Targeted partial reprogramming of age-associated cell states improves markers of health in mouse models of aging Sci Transl Med 16, eadg1777 10.1126/scitranslmed.adg1777

33. Parras, A., Vílchez-Acosta, A., Desdín-Micó, G., Picó, S., Mrabti, C., Montenegro-Borbolla, E. et al. (2023) In vivo reprogramming leads to premature death linked to hepatic and intestinal failure Nature Aging 3, 1509-1520 10.1038/s43587-023-00528-5

34. Lu, Y., Brommer, B., Tian, X., Krishnan, A., Meer, M., Wang, C. et al. (2020) Reprogramming to recover youthful epigenetic information and restore vision Nature 588, 124–129 10.1038/s41586-020-2975-4

35. Macip, C. C., Hasan, R., Hoznek, V., Kim, J., Lu, Y. R., Metzger, L. E. t., et al. (2024) Gene Therapy-Mediated Partial Reprogramming Extends Lifespan and Reverses Age-Related Changes in Aged Mice Cell Reprogram 26, 24–32 10.1089/cell.2023.0072

36. Ksander, B., Shah, M., Krasniqi, D., Gregory-Ksander, M. S., Rosenzweig-Lipson, S., Broniowska, K. et al. (2023) Epigenetic reprogramming-A novel gene therapy that restores vision loss in a nonhuman primate model of NAION Investigative Ophthalmology & Visual Science 64, 474–474,

37. Pereira, B., Correia, F. P., Alves, I. A., Costa, M., Gameiro, M., Martins, A. P. et al. (2024) Epigenetic reprogramming as a key to reverse ageing and increase longevity Ageing Research Reviews 95, 102204 10.1016/j.arr.2024.102204

38. Liuyang, S., Wang, G., Wang, Y., He, H., Lyu, Y., Cheng, L. et al. (2023) Highly efficient and rapid generation of human pluripotent stem cells by chemical reprogramming Cell Stem Cell 30, 450–459.e459 10.1016/j.stem.2023.02.008

39. Wang, Y., Peng, F., Yang, Z., Cheng, L., Cao, J., Fu, X., et al. (2025) A rapid chemical reprogramming system to generate human pluripotent stem cells Nature Chemical Biology 10.1038/s41589-024-01799-8

40. Guan, J., Wang, G., Wang, J., Zhang, Z., Fu, Y., Cheng, L. et al. (2022) Chemical reprogramming of human somatic cells to pluripotent stem cells Nature 605, 325–331 10.1038/s41586-022-04593-5

41. Wang, S., Du, Y., Zhang, B., Meng, G., Liu, Z., Liew, S. Y. et al. (2024) Transplantation of chemically induced pluripotent stem-cell-derived islets under abdominal anterior rectus sheath in a type 1 diabetes patient Cell 187, 6152-6164.e6118 10.1016/j.cell.2024.09.004

42. Schoenfeldt, L., Paine, P. T., Kamaludeen M, N. H., Phelps, G. B., Mrabti, C., Perez, K. et al. (2022) Chemical reprogramming ameliorates cellular hallmarks of aging and extends lifespan bioRxiv 2022.2008.2029.505222 10.1101/2022.08.29.505222

43. Mitchell, W., Goeminne, L. J. E., Tyshkovskiy, A., Zhang, S., Chen, J. Y., Paulo, J. A., et al. (2024) Multi-omics characterization of partial chemical reprogramming reveals evidence of cell rejuvenation Elife 12, 10.7554/eLife.90579

44. Yang, J. H., Petty, C. A., Dixon-McDougall, T., Lopez, M. V., Tyshkovskiy, A., Maybury-Lewis, S. et al. (2023) Chemically induced reprogramming to reverse cellular aging Aging (Albany NY) 15, 5966-5989 10.18632/aging.204896

45. Schindelin, J., Arganda-Carreras, I., Frise, E., Kaynig, V., Longair, M., Pietzsch, T. et al. (2012) Fiji: an open-source platform for biological-image analysis Nature Methods **9**, 676-682 10.1038/nmeth.2019

46. Chaudhry, A., Shi, R., and Luciani, D. S. (2020) A pipeline for multidimensional confocal analysis of mitochondrial morphology, function, and dynamics in pancreatic β-cells Am J Physiol Endocrinol Metab 318, E87-e101 10.1152/ajpendo.00457.2019

47. Lefebvre, A., Ma, D., Kessenbrock, K., Lawson, D. A., and Digman, M. A. (2021) Automated segmentation and tracking of mitochondria in live-cell time-lapse images Nat Methods 18, 1091-1102 10.1038/s41592-021-01234-z

48. Arganda-Carreras, I., Kaynig, V., Rueden, C., Eliceiri, K. W., Schindelin, J., Cardona, A. et al. (2017) Trainable Weka Segmentation: a machine learning tool for microscopy pixel classification Bioinformatics 33, 2424-2426 10.1093/bioinformatics/btx180

49. Bankhead, P., Loughrey, M. B., Fernández, J. A., Dombrowski, Y., McArt, D. G., Dunne, P. D., et al. (2017) QuPath: Open source software for digital pathology image analysis Scientific Reports 7, 16878 10.1038/s41598-017-17204-5

50. Mitchell, W., Pharaoh, G., Tyshkovskiy, A., Campbell, M., Marcinek, D. J., and Gladyshev, V. N. (2025) The Mitochondria-Targeted Peptide Therapeutic Elamipretide Improves Cardiac and Skeletal Muscle Function During Aging Without Detectable Changes in Tissue Epigenetic or Transcriptomic Age Aging Cell n/a, e70026 10.1111/acel.70026

51. Robinson, M. D., McCarthy, D. J., and Smyth, G. K. (2010) edgeR: a Bioconductor package for differential expression analysis of digital gene expression data Bioinformatics 26, 139–140 10.1093/bioinformatics/btp616

52. Tyshkovskiy, A., Ma, S., Shindyapina, A. V., Tikhonov, S., Lee, S. G., Bozaykut, P. et al. (2023) Distinct longevity mechanisms across and within species and their association with aging Cell 186, 2929-2949.e2920 10.1016/j.cell.2023.05.002

53. Kriukov, D., Khrameeva, E. E., Gladyshev, V. N., Dmitriev, S. E., and Tyshkovskiy, A. (2022) Longevity and rejuvenation effects of cell reprogramming are decoupled from loss of somatic identity bioRxiv 2022.2012.2012.520058 10.1101/2022.12.12.520058

54. Goeminne, L. J., Gevaert, K., and Clement, L. (2016) Peptide-level Robust Ridge Regression Improves Estimation, Sensitivity, and Specificity in Data-dependent Quantitative Label-free Shotgun Proteomics Mol Cell Proteomics 15, 657–668 10.1074/mcp.M115.055897

55. Goeminne, L. J. E., Gevaert, K., and Clement, L. (2018) Experimental design and data-analysis in label-free quantitative LC/MS proteomics: A tutorial with MSqRob J Proteomics 171, 23–36 10.1016/j.jprot.2017.04.004

56. Benjamini, Y., and Hochberg, Y. (1995) Controlling the False Discovery Rate: A Practical and Powerful Approach to Multiple Testing Journal of the Royal Statistical Society Series B (Methodological) 57, 289–300, http://www.jstor.org/stable/2346101

57. Ritchie, M. E., Phipson, B., Wu, D., Hu, Y., Law, C. W., Shi, W., et al. (2015) limma powers differential expression analyses for RNA-sequencing and microarray studies Nucleic Acids Research **43**, e47-e47 10.1093/nar/gkv007

58. Baker, N., Patel, J., and Khacho, M. (2019) Linking mitochondrial dynamics, cristae remodeling and supercomplex formation: How mitochondrial structure can regulate bioenergetics Mitochondrion 49, 259–268 10.1016/j.mito.2019.06.003

59. Worthley, E. G., and Schott, C. D. (1969) The toxicity of four concentrations of DMSO Toxicology and Applied Pharmacology 15, 275–281 10.1016/0041-008X(69)90027-1

60. Galvao, J., Davis, B., Tilley, M., Normando, E., Duchen, M. R., and Cordeiro, M. F. (2014) Unexpected low-dose toxicity of the universal solvent DMSO Faseb j 28, 1317-1330 10.1096/fj.13-235440

61. Sweetwyne, M. T., Pippin, J. W., Eng, D. G., Hudkins, K. L., Chiao, Y. A., Campbell, M. D. et al. (2017) The mitochondrial-targeted peptide, SS-31, improves glomerular architecture in mice of advanced age Kidney International **91**, 1126-1145 10.1016/j.kint.2016.10.036

62. Hennigar, R. A., Schulte, B. A., and Spicer, S. S. (1985) Heterogeneous distribution of glycoconjugates in human kidney tubules Anat Rec 211, 376–390 10.1002/ar.1092110403

63. Ichimura, T., Bonventre, J. V., Bailly, V., Wei, H., Hession, C. A., Cate, R. L. et al. (1998) Kidney injury molecule-1 (KIM-1), a putative epithelial cell adhesion molecule containing a novel immunoglobulin domain, is up-regulated in renal cells after injury J Biol Chem 273, 4135-4142 10.1074/jbc.273.7.4135

64. Kellerman, P. S., and Bogusky, R. T. (1992) Microfilament disruption occurs very early in ischemic proximal tubule cell injury Kidney Int 42, 896–902 10.1038/ki.1992.366

65. Takahashi, N., Boysen, G., Li, F., Li, Y., and Swenberg, J. A. (2007) Tandem mass spectrometry measurements of creatinine in mouse plasma and urine for determining glomerular filtration rate Kidney International 71, 266–271 10.1038/sj.ki.5002033

66. El-Zayadi, A. R. (2008) Hepatic steatosis: a benign disease or a silent killer World J Gastroenterol 14, 4120-4126 10.3748/wjg.14.4120

67. Smith, Z. D., Sindhu, C., and Meissner, A. (2016) Molecular features of cellular reprogramming and development Nature Reviews Molecular Cell Biology 17, 139–154 10.1038/nrm.2016.6

68. Zhao, T., Fu, Y., Zhu, J., Liu, Y., Zhang, Q., Yi, Z. et al. (2018) Single-Cell RNA-Seq Reveals Dynamic Early Embryonic-like Programs during Chemical Reprogramming Cell Stem Cell 23, 31–45.e37 10.1016/j.stem.2018.05.025

69. Fan, H., and Tan, Y. (2024) Lipid Droplet-Mitochondria Contacts in Health and Disease Int J Mol Sci 25, 10.3390/ijms25136878

70. Cui, L., and Liu, P. (2020) Two Types of Contact Between Lipid Droplets and Mitochondria Frontiers in Cell and Developmental Biology Volume 8 - 2020, 10.3389/fcell.2020.618322

71. Talari, N. K., Mattam, U., Meher, N. K., Paripati, A. K., Mahadev, K., Krishnamoorthy, T., et al. (2023) Lipid-droplet associated mitochondria promote fatty-acid oxidation through a distinct bioenergetic pattern in male Wistar rats Nature Communications 14, 766 10.1038/s41467-023-36432-0

