## Supporting Information for "*In vivo* chemical reprogramming is associated with a toxic accumulation of lipid droplets hindering rejuvenation"

**Table of contents:**

**Table S1** – Drug supplier and dosage information

**Table S2** – Liver differentially expressed genes

**Table S3** – Kidney differentially expressed genes

**Table S4** – Significant liver GSEA terms

**Table S5** – Significant kidney GSEA terms

**Supplemental Figure S1** – Effect of uncoupler treatment on mitochondrial morphology and networks

**Supplemental Figure S2** – Effect of *in vivo* partial chemical reprogramming on liver and kidney tissue transcriptomic age

**Supplemental Figure S3** – Additional kidney histology images

**Supplemental Figure S4** – Modular transcriptomic age of fibroblasts treated with 2c and 7c

**Table S1: Drug supplier and dosage information**

| Compound | Supplier | Catalog # | Cell Culture Concentration (μM) | Trial 1 Dose (mg/kg/day) | Trial 2 Dose (mg/kg/day) | Trial 3 Dose (mg/kg/day) |
|---|---|---|---|---|---|---|
| Valproate | STEMCELL Technologies | 72292 | 500 | 1 | 50 | 10 |
| Repsox | Sigma-Aldrich | R0158 | 5 | 0.1 | 0.5 | 0.25 |
| Tranylcypromine | Sigma-Aldrich | P22370 | 5 | 0.1 | 0.5 | 0.25 |
| Forskolin | Tocris | 1099 | 10 | 0.1 | 0.5 | 0.25 |
| CHIR99021 | Cayman Chemical | 13122 | 10 | 0.1 | 0.5 | 0.25 |
| DZNep | APExBIO | A1905 | 0.5 | 0.1 | 0.5 | 0.25 |
| TTNPB | Cayman Chemical | 16144 | 1 | 0.1 | 0.5 | 0.25 |

### Table S1

**Table S2: Liver differentially expressed genes**

| Entrez ID | Gene Symbol | log2FC | P-value | adjusted P-value |
|---|---|---|---|---|
| 71640 | Zfp949 | -1.282 | 5.94E-14 | 7.82E-10 |
| 100040633 | Rpsa-ps10 | -3.240 | 4.83E-08 | 3.18E-04 |
| 13116 | Cyp46a1 | 4.259 | 5.25E-07 | 2.30E-03 |
| 116904 | Alpk3 | 4.536 | 1.53E-06 | 4.57E-03 |
| 214305 | Hhipl1 | 2.443 | 1.74E-06 | 4.57E-03 |
| 22153 | Tubb4a | 3.971 | 3.40E-06 | 7.45E-03 |
| 228852 | Ppp1r16b | -1.073 | 1.49E-05 | 2.80E-02 |

### Table S2

**Table S3: Kidney differentially expressed genes**

| Entrez ID | Gene Symbol | log2FC | P-value | adjusted P-value |
|---|---|---|---|---|
| 244958 | Mrap2 | -3.274 | 1.11E-13 | 1.67E-09 |
| 71640 | Zfp949 | -1.025 | 3.78E-11 | 2.85E-07 |
| 100040633 | Rpsa-ps10 | -2.551 | 4.84E-10 | 2.43E-06 |
| 13884 | Ces1c | -2.529 | 4.40E-09 | 1.66E-05 |
| 17842 | Mup3 | -2.771 | 5.76E-07 | 1.74E-03 |
| 21835 | Thrsp | 2.448 | 9.15E-07 | 2.30E-03 |
| 98660 | Atp1a2 | 1.077 | 2.58E-06 | 5.56E-03 |
| 13095 | Cyp2c29 | -2.600 | 1.78E-05 | 3.16E-02 |
| 14061 | F2 | -2.108 | 1.88E-05 | 3.16E-02 |
| 14311 | Cidec | 2.538 | 2.39E-05 | 3.61E-02 |

### Table S3

**Table S4: Significant liver GSEA terms**

| Term | NES | P-value | adjusted P-value |
|---|---|---|---|
| REACTOME SRP DEPENDENT COTRANSLATIONAL PROTEIN TARGETING TO MEMBRANE | 2.193 | 2.84E-08 | 2.63E-05 |
| REACTOME RESPIRATORY ELECTRON TRANSPORT | 2.187 | 4.68E-08 | 2.63E-05 |
| REACTOME RESPIRATORY ELECTRON TRANSPORT ATP SYNTHESIS BY CHEMIOSMOTIC COUPLING AND HEAT PRODUCTION BY UNCOUPLING PROTEINS | 2.173 | 7.43E-08 | 2.78E-05 |
| REACTOME CELLULAR RESPONSE TO STARVATION | 2.019 | 1.70E-07 | 4.76E-05 |
| REACTOME SELENOAMINO ACID METABOLISM | 2.141 | 4.04E-07 | 9.07E-05 |
| REACTOME RESPONSE OF EIF2AK4 GCN2 TO AMINO ACID DEFICIENCY | 2.166 | 4.90E-07 | 9.16E-05 |
| KEGG RIBOSOME | 2.183 | 6.70E-07 | 9.61E-05 |
| REACTOME THE CITRIC ACID TCA CYCLE AND RESPIRATORY ELECTRON TRANSPORT | 1.975 | 6.85E-07 | 9.61E-05 |
| REACTOME METABOLISM OF AMINO ACIDS AND DERIVATIVES | 1.763 | 7.89E-07 | 9.85E-05 |
| REACTOME EUKARYOTIC TRANSLATION ELONGATION | 2.130 | 2.41E-06 | 2.47E-04 |
| REACTOME TRANSLATION | 1.778 | 2.47E-06 | 2.47E-04 |
| HALLMARK OXIDATIVE PHOSPHORYLATION | 1.894 | 2.64E-06 | 2.47E-04 |
| REACTOME REGULATION OF EXPRESSION OF SLITS AND ROBOS | 1.846 | 1.10E-05 | 9.51E-04 |
| KEGG OXIDATIVE PHOSPHORYLATION | 2.018 | 2.31E-05 | 1.85E-03 |
| REACTOME SELECTIVE AUTOPHAGY | 1.941 | 6.00E-05 | 4.49E-03 |
| REACTOME COMPLEX I BIOGENESIS | 1.986 | 8.28E-05 | 5.81E-03 |
| REACTOME INFLUENZA INFECTION | 1.764 | 9.28E-05 | 6.13E-03 |
| REACTOME AGGREPHAGY | 2.064 | 1.35E-04 | 8.39E-03 |
| REACTOME MHC CLASS II ANTIGEN PRESENTATION | 1.809 | 1.42E-04 | 8.39E-03 |
| REACTOME HSP90 CHAPERONE CYCLE FOR STEROID HORMONE RECEPTORS SHR IN THE PRESENCE OF LIGAND | 1.977 | 1.88E-04 | 1.05E-02 |
| HALLMARK ALLOGRAFT REJECTION | 1.701 | 2.50E-04 | 1.22E-02 |
| KEGG PARKINSONS DISEASE | 1.890 | 2.34E-04 | 1.22E-02 |
| KEGG PRIMARY IMMUNODEFICIENCY | 1.983 | 2.46E-04 | 1.22E-02 |
| REACTOME NONSENSE MEDIATED DECAY NMD | 1.821 | 2.69E-04 | 1.26E-02 |
| REACTOME GAP JUNCTION ASSEMBLY | 1.925 | 3.37E-04 | 1.51E-02 |
| REACTOME SIGNALING BY ROBO RECEPTORS | 1.660 | 3.49E-04 | 1.51E-02 |
| KEGG ANTIGEN PROCESSING AND PRESENTATION | 1.892 | 3.75E-04 | 1.56E-02 |
| REACTOME ROS AND RNS PRODUCTION IN PHAGOCYTES | 1.952 | 4.01E-04 | 1.61E-02 |
| REACTOME EUKARYOTIC TRANSLATION INITIATION | 1.790 | 4.17E-04 | 1.62E-02 |
| REACTOME AUTOPHAGY | 1.641 | 6.42E-04 | 2.40E-02 |
| REACTOME CELLULAR RESPONSE TO CHEMICAL STRESS | 1.650 | 6.62E-04 | 2.40E-02 |
| HALLMARK ADIPOGENESIS | 1.614 | 7.84E-04 | 2.70E-02 |
| REACTOME POST CHAPERONIN TUBULIN FOLDING PATHWAY | 1.870 | 7.93E-04 | 2.70E-02 |
| REACTOME THE ROLE OF GTSE1 IN G2 M PROGRESSION AFTER G2 CHECKPOINT | 1.762 | 9.42E-04 | 3.02E-02 |
| KEGG SYSTEMIC LUPUS ERYTHEMATOSUS | 1.822 | 9.22E-04 | 3.02E-02 |
| KEGG GLYCOLYSIS GLUCONEOGENESIS | 1.833 | 1.02E-03 | 3.19E-02 |
| REACTOME CYTOPROTECTION BY HMOX1 | 1.630 | 1.16E-03 | 3.43E-02 |
| KEGG HUNTINGTONS DISEASE | 1.627 | 1.16E-03 | 3.43E-02 |

### Table S4

**Table S5: Significant kidney GSEA terms**

| Term | NES | P-value | adjusted P-value |
|---|---|---|---|
| KEGG RIBOSOME | 1.979 | 1.50E-05 | 8.36E-03 |
| HALLMARK MYOGENESIS | 1.904 | 3.38E-06 | 3.91E-03 |
| REACTOME EUKARYOTIC TRANSLATION ELONGATION | 1.992 | 2.17E-05 | 8.36E-03 |
| KEGG LINOLEIC ACID METABOLISM | -2.020 | 4.69E-05 | 1.35E-02 |

### Table S5

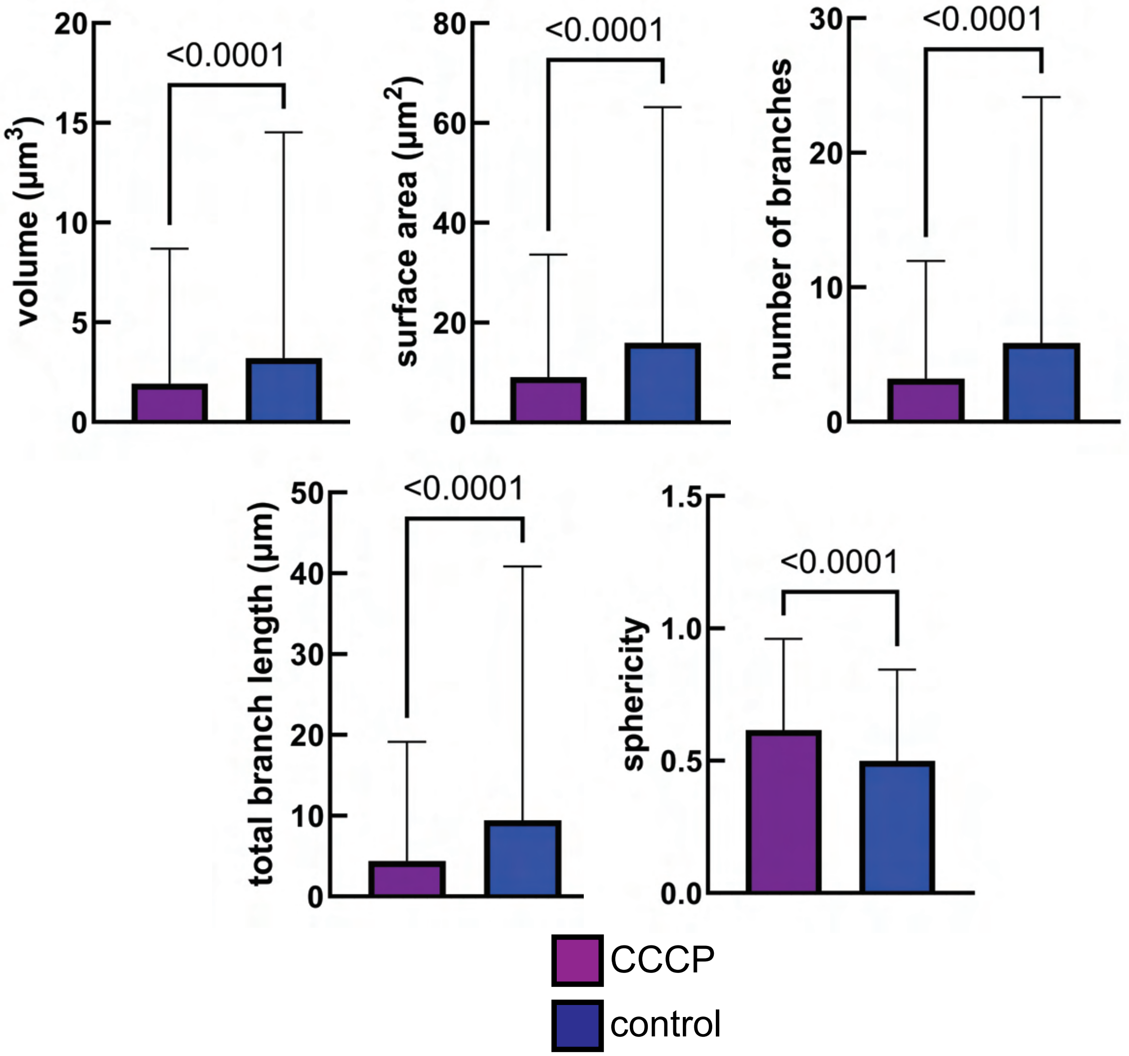


### Supplemental Figure S1

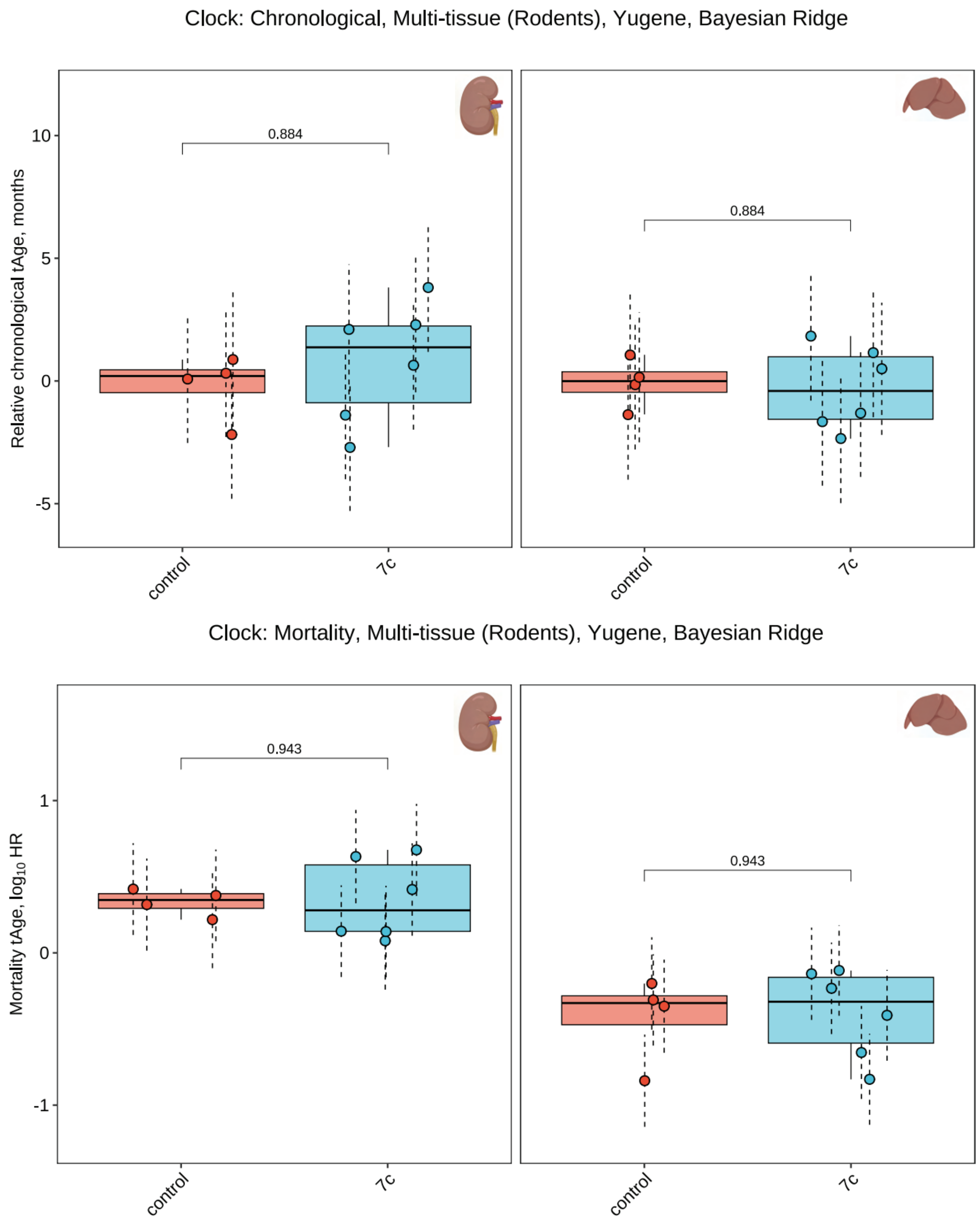


### Supplemental Figure S2

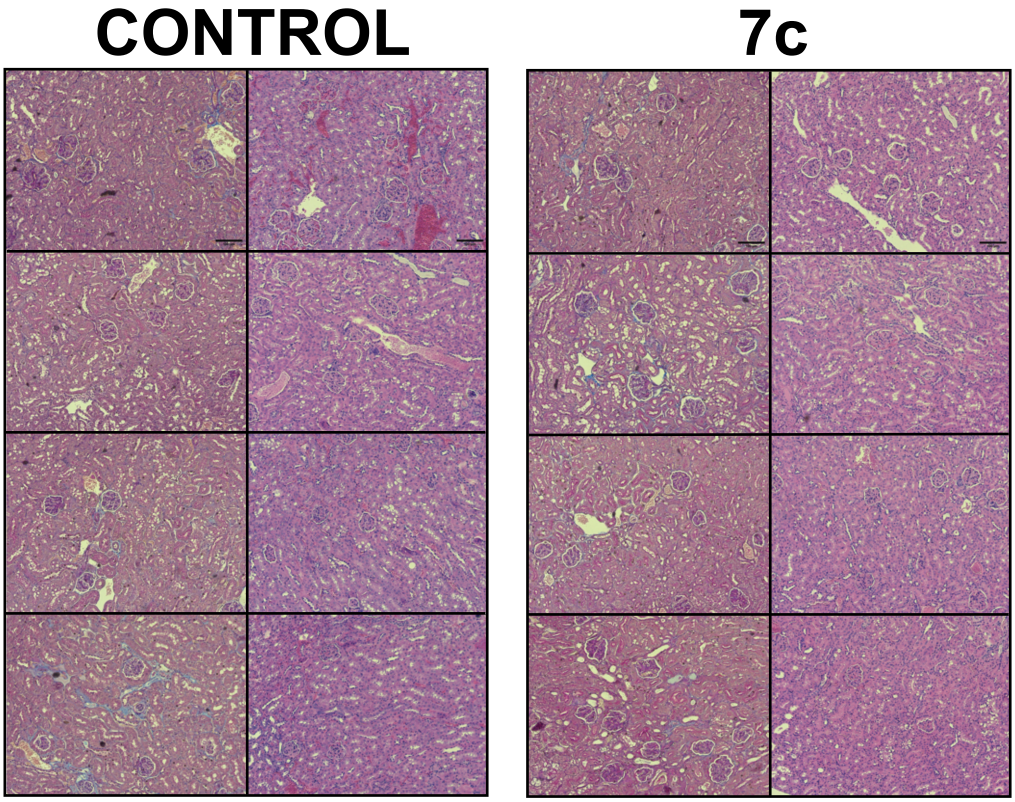


### Supplemental Figure S3

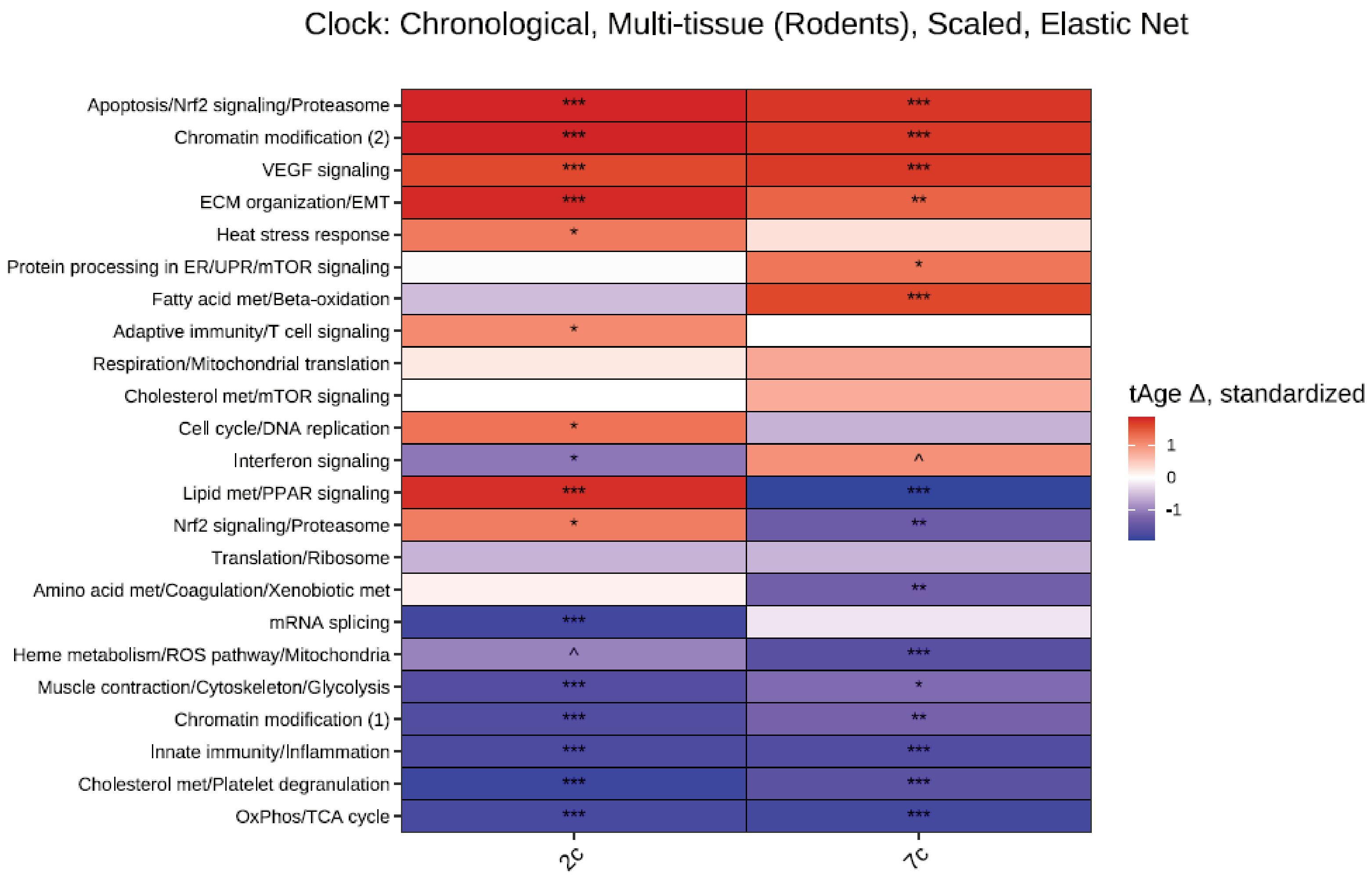


### Supplemental Figure S4
